# Conditioned pain modulation recruits the descending pain modulatory system to inhibit spinal activity

**DOI:** 10.64898/2026.03.10.710789

**Authors:** Alexandra Tinnermann, Karita E. Ojala, Björn Horing, Ying Chu, Jürgen Finsterbusch, Christian Büchel

**Author notes:** **Corresponding author:** Alexandra Tinnermann. These authors contributed equally (Alexandra Tinnermann, Karita E. Ojala, Björn Horing).

## Abstract

Conditioned pain modulation (CPM) is a phenomenon where the perception of pain is reduced in the presence of a second painful stimulus. In rodents, a similar phenomenon involves descending control pathways that rely on the brainstem to modulate spinal cord activity while in humans, descending mechanisms including the spinal cord remain elusive. In this study, we applied functional MRI to simultaneously record brain and spinal cord responses during CPM in healthy participants. CPM developed over time and was associated with reduced activity in key pain-related brain regions and the spinal cord dorsal horn. Conversely, regions involved in descending pain modulation showed increased activity during CPM. Functional connectivity analyses revealed reduced coupling during CPM between spinal and descending control regions. Our findings demonstrate that CPM involves both the descending pain control system to inhibit spinal activity and bottom-up pain signalling of attenuated spinal responses.

## Introduction

Pain serves as a vital warning system, alerting us to potential or actual tissue damage with the goal to avoid harm. While experiencing pain is necessary for survival, the nervous system also possesses endogenous mechanisms to modulate pain perception depending on the context. For example, in situations that are not immediately life-threatening, persistent pain can be inhibited if a more relevant threat emerges or if the organism needs to focus on other tasks (Fields, 2004; Seymour, 2019). A key component of such pain modulation involves descending pathways originating from cortical areas like the ventromedial prefrontal cortex (vmPFC) and anterior cingulate cortex (ACC) which project to brainstem nuclei, including the periaqueductal gray (PAG) and rostral ventromedial medulla (RVM), descending further to modulate pain transmission in the spinal cord. Within the spinal cord, descending signals can inhibit the activity of neurons that transmit pain signals to the brain, effectively reducing pain perception (Bannister, 2019; Fields, 2004; Sirucek et al., 2023).

A well-studied example of this endogenous pain modulation is the "pain inhibits pain" phenomenon, where painful stimuli applied at one body site can diminish pain perception at another body site. In animals, this mechanism is known as Diffuse Noxious Inhibitory Controls (DNIC), which is mediated by brainstem systems (Bars et al., 1979; Kucharczyk et al., 2023, 2022; Villanueva and Le Bars, 1995) thus suggesting the involvement of the descending pain modulatory system.

In humans, a perceptually similar phenomenon known as conditioned pain modulation (CPM) can be experimentally induced by applying one painful stimulus to reduce the perception of another painful stimulus at a different body site. The efficacy of CPM varies among individuals and has been proposed as a marker of descending control efficacy, which might be an important factor for the vulnerability for pain chronification (Yarnitsky et al., 2008). Accordingly, individuals with chronic pain states such as fibromyalgia, migraine, irritable bowel syndrome or osteoarthritis show impaired CPM (Arendt-Nielsen et al., 2010; Julien et al., 2005; Kosek and Hansson, 1997; Piché et al., 2010; Pielsticker et al., 2005; Sandrini et al., 2006). Despite its clinical relevance, the neural mechanisms underlying CPM in humans remain only partially understood (Sirucek et al., 2023). Neuroimaging studies investigating CPM have identified decreased activity in pain-related brain regions such as the thalamus, insula, secondary somatosensory cortex and mid-cingulate cortex (Bogdanov et al., 2015; Kisler et al., 2018; Nahman-Averbuch et al., 2014; Piché et al., 2009; Song et al., 2006; Sprenger et al., 2011). Furthermore, CPM-related analgesia was correlated with activity in different brainstem nuclei (Youssef et al., 2016a) and functional connectivity between the subgenual ACC, PAG and RVM (Li et al., 2025; Sprenger et al., 2011). Notably, the role of the spinal cord as the primary entry point of nociceptive signals has not been directly examined in humans during CPM, despite its central role in the pain modulation network (Eippert et al., 2009b; Sprenger et al., 2012; Tinnermann et al., 2017). Therefore, in this study we simultaneously acquired fMRI data in the brain, brainstem, and cervical spinal cord (Finsterbusch et al., 2013; Tinnermann et al., 2021) while healthy participants underwent a CPM paradigm using pressure pain. Tonic painful stimuli (conditioning stimuli) were applied to one arm while phasic painful stimuli (test stimuli) were applied to the other arm (Fig. 1). After every phasic stimulus, participants rated the painfulness of the stimulus on a Visual Analogue Scale (VAS). As a control condition, we implemented tonic non-painful stimuli paired with phasic painful stimuli to account for unspecific attention and sensory stimulation effects. This allowed us to characterize CPM in the entire central nervous system.

**Figure 1.**
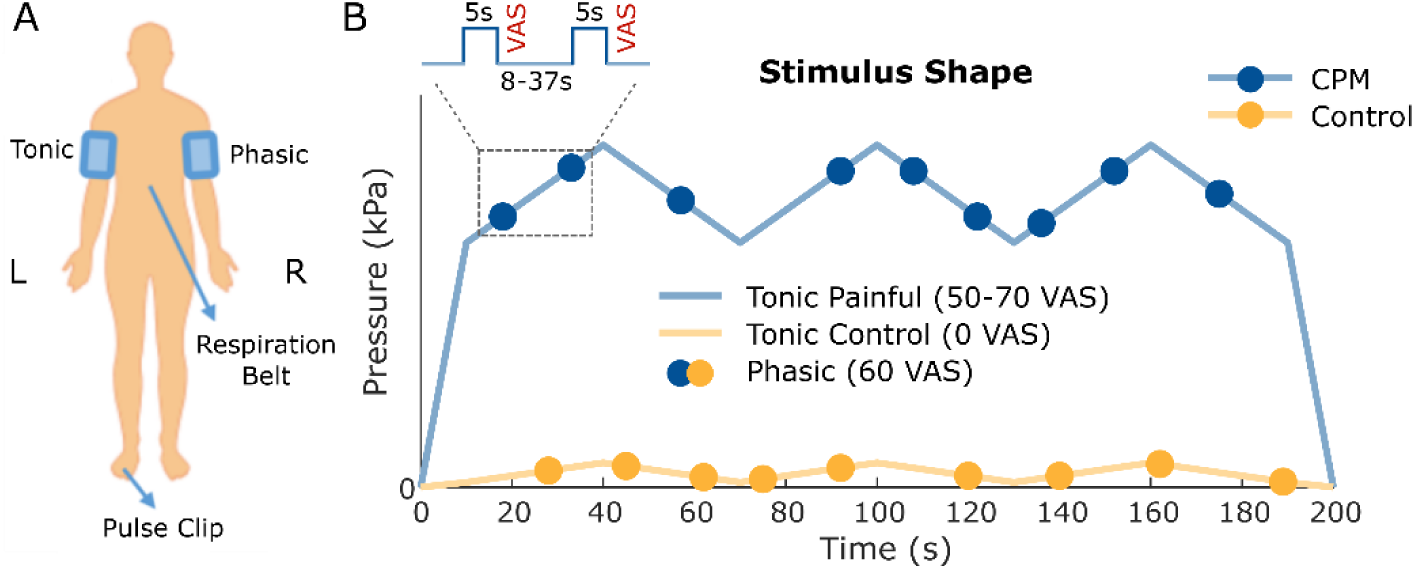
Experimental design. (**A**) Tonic stimuli were applied on the left arm and phasic stimuli on the right arm via automatized pressure cuff algometry. Respiration was measured with a belt around the chest and pulse was measured on the left big toe. (**B**) Painful tonic stimuli fluctuated slightly, completing three cycles within one stimulus with a total duration of 200 s. The control tonic stimuli also fluctuated slightly with very low pressure output. Nine phasic stimuli of 5 s duration (dots) were randomly distributed along predetermined time points in the tonic stimulus with three stimuli per cycle (interstimulus interval 8 to 37 s). After every phasic stimulus, participants rated their pain intensity on a VAS scale. In total, participants received 72 phasic stimuli across four fMRI runs with 36 stimuli per condition.

## Results

### Behavioural results

In a first step, we calculated the mean phasic pain ratings for CPM and control conditions (Fig. 2A). 45% of participants (N=19) reported CPM-related analgesia, around 10% of participants (N=4) showed no difference between CPM and control ratings while 45% of participants (N=19) reported hyperalgesia (Fig. 2B). Across all participants, there was no significant difference between CPM and control ratings (ΔVAS = 0.92, t_41_ = 0.64, p = 0.53). Given that the aversiveness of the tonic stimulus increased over time (Fig. S1A), we next investigated whether CPM effects varied over time. This analysis revealed a significant interaction between CPM versus control condition and trial number (mean±SE: −0.109±0.028, F[1,2943] = 15.4, p = 0.00008) in a way that the CPM effect increased over time (Fig. 2C). In addition, previous studies have shown that CPM depends on the painfulness of the stimuli with stronger CPM for more painful stimuli (Chaudhry et al., 2025; Nir et al., 2011). Therefore, we tested whether the individual painfulness of the tonic stimulus (Fig. S1B) contributed to CPM. This analysis showed a significant interaction between CPM versus control condition and the tonic rating provided at the end of the experiment (mean±SE: −0.065±0.026, F[1,2943] = 6.16, p = 0.013), indicating that participants who rated the tonic stimulus as more painful also experienced a stronger CPM effect.

**Figure 2.**
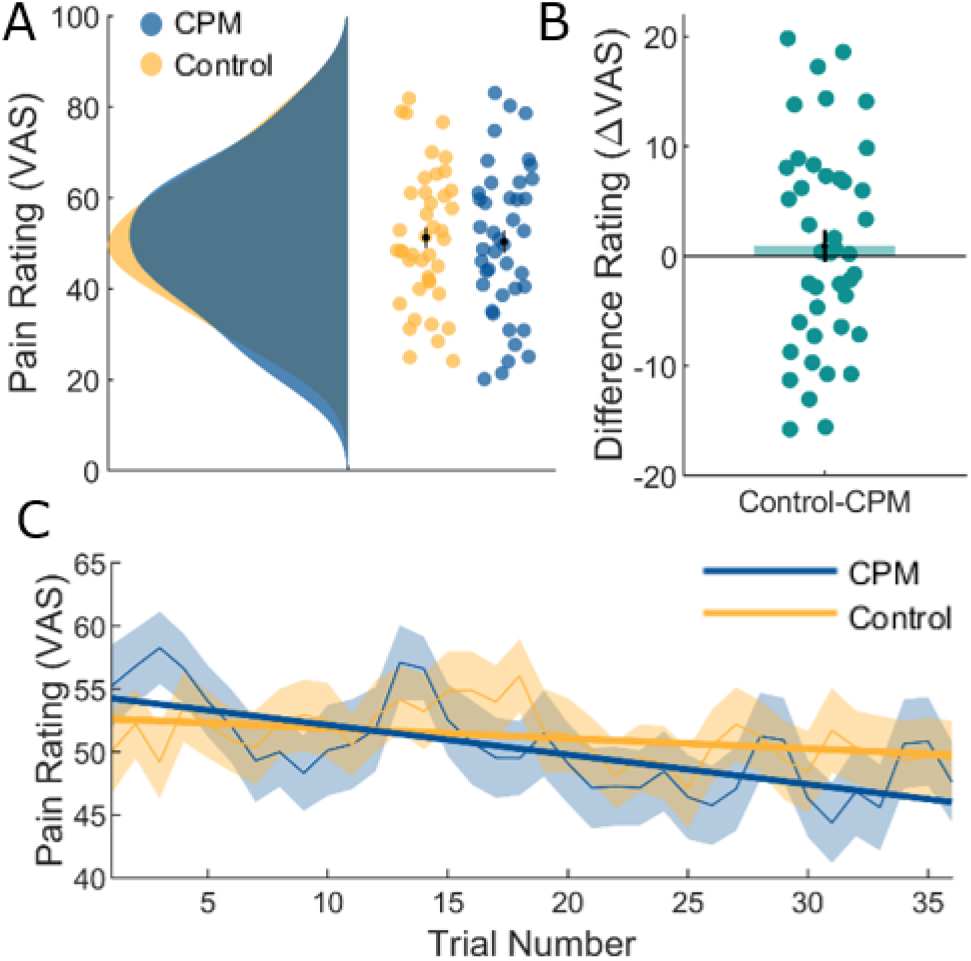
Behavioural results. (**A**) Mean phasic pain ratings for CPM and control. CPM and control ratings did not differ significantly across participants. (**B**) Distribution of difference ratings between control and CPM conditions. (**C**) Development of CPM over all 36 phasic stimuli compared to control stimuli. Solid thick lines show a linear fit separately for control and CPM. The significant decrease in pain ratings over time was more pronounced in the CPM condition compared to the control condition (interaction effect p < 0.001). Please note that this visualization does not fully reflect the implemented linear mixed model as with our fully counterbalanced design, condition order cannot be integrated.

### Pressure-related BOLD responses across brain and spinal cord

As a sanity check, we analysed the main effect of phasic stimulation in the brain and spinal cord. Typical pain-processing regions such as the operculum, insula, thalamus, cingulate cortex, pre- and postcentral gyrus showed increased BOLD responses during phasic pressure cuff stimulation (Fig. S2A). Moreover, the right dorsal horn ipsilateral to phasic stimulation site showed increased activity at vertebral level C5 which corresponds to spinal segment C6 (Fig. S2B). For the subsequent analyses, we mainly report analyses investigating the time course of brain and spinal cord responses since a previous study observed that pressure pain elicited a pronounced stimulus-onset response which quickly decayed (Nold et al., 2025b). Therefore, we conducted first-level models based on Finite Impulse Response (FIR) functions allowing us to look at post-stimulus averaged fMRI signal over time with the advantage of not being constrained by a particular hemodynamic response function. Response functions in the parietal and frontal operculum, the insula, postcentral gyrus, thalamus and brainstem regions such as the PAG and RVM and the spinal cord dorsal horn are shown in Figure S3. Temporal profiles across regions indicate two distinct response patterns. First, BOLD responses in regions such as the parietal operculum (SII), posterior insula, and RVM were quite short and other regions such as the pre-/postcentral gyrus (SI), anterior insula, thalamus and operculum showed a bimodal response with a second peak after stimulus offset resulting in a prolonged signal.

### BOLD differences between CPM and the control condition across brain and spinal cord

First, we analysed which regions showed decreased activity during the CPM condition compared to control therefore reflecting CPM effects. This analysis revealed decreased activation during the CPM condition in the frontal operculum (xyz_MNI_: 50/0/6, t_760_ = 8.21, p_SVC_ < 0.001), anterior insula (xyz_MNI_: 38/10/9, t_760_ = 7.70, p_SVC_ < 0.001), mid-cingulate cortex (xyz_MNI_: −12/12/34, t_760_ = 5.63, p_SVC_ < 0.001), posterior insula (xyz_MNI_: −39/-10/3, t_760_ = 5.63, p_SVC_ < 0.001), parietal operculum (xyz_MNI_: 51/-20/16, t_760_ = 5.51, p_SVC_ < 0.001), and thalamus (xyz_MNI_: 4/-12/12, t_760_ = 4.88, p_SVC_ = 0.007; Fig. 3).

**Figure 3.**
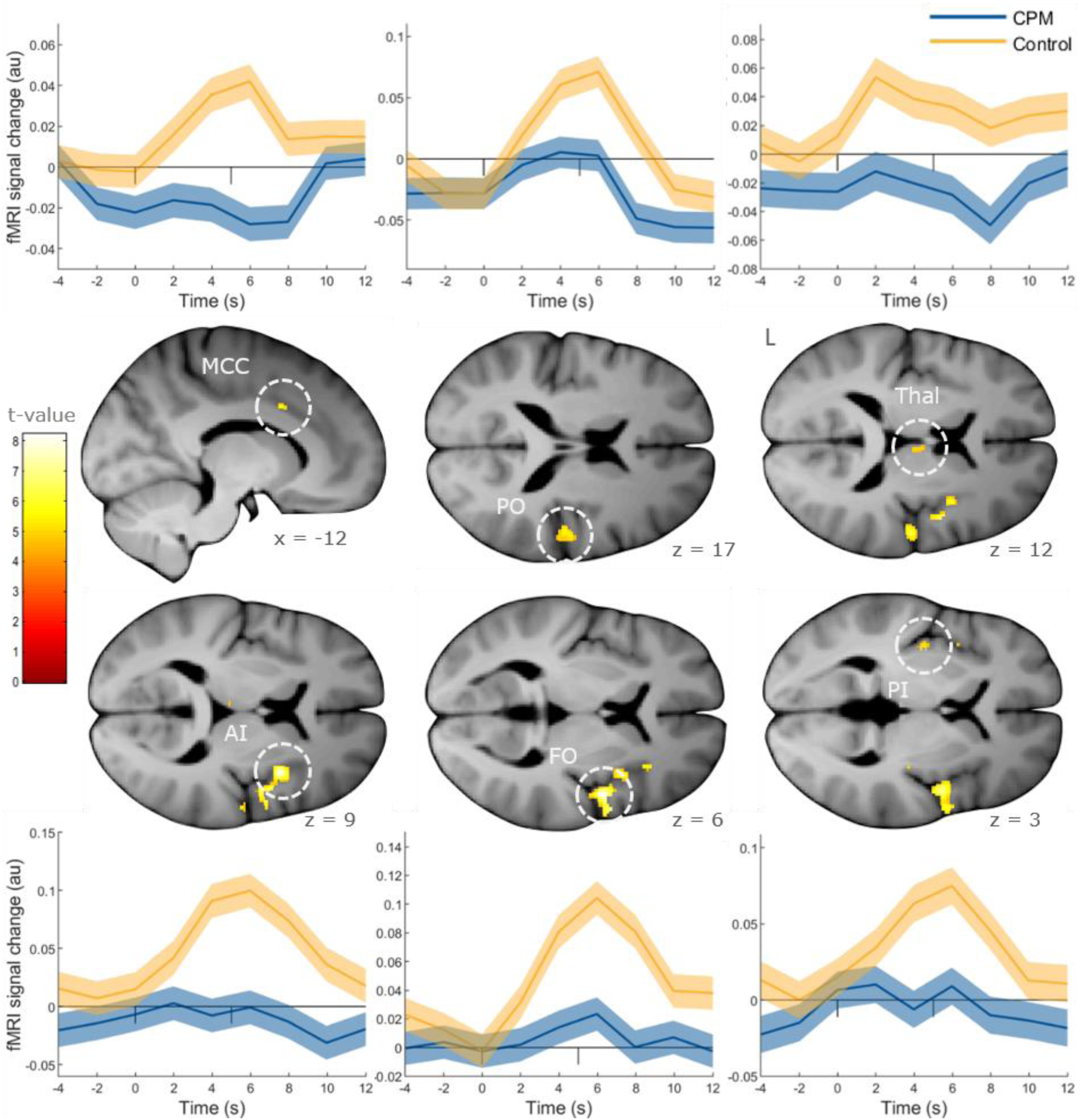
Pain-related brain regions showing CPM effects in a FIR model. Time courses in opercular and insular regions as well as in the cingulate cortex and thalamus show reduced activation in the CPM condition compared to control. Pain-related region-of-interest masked statistical t-maps are overlaid on an average structural T1 image in Montreal Neurological Institute (MNI) template space and the visualization threshold is set to p_SVC_ < 0.05. Dashed circles indicate the cluster from which time courses of peak voxel parameter estimates are plotted. Shaded areas represent standard error of the mean. Ticks on the zero line indicate stimulus duration. Please note that the BOLD signal at a given time point reflects the averaged BOLD signal within one TR starting at that time point. MCC: mid-cingulate cortex, PO: parietal operculum, Thal: thalamus, AI: anterior insula, FO: frontal operculum, PI: posterior insula.

With regard to the brainstem, the RVM (xyz_MNI_: 2/-33/-46, t_760_ = 4.59, p_SVC_ = 0.024, Fig 4A) showed reduced activity in the CPM condition compared to the control condition. Moreover, two clusters in the right dorsal horn of the spinal cord at different heights of spinal segment C6 showed reduced activity in the CPM condition (xyz_MNI_: 4/-49/-142, t = 3.51, p_SVC_ = 0.015; xyz_MNI_: 3/-48/-155, t = 3.29, p_SVC_ = 0.030, Fig. 4B). Both clusters were located ipsilateral to the stimulation site and at the height of the upper and lower disk spanning vertebrae C5, respectively. Individual CPM effect magnitudes did not correlate with BOLD responses in the brain or spinal cord.

**Figure 4.**
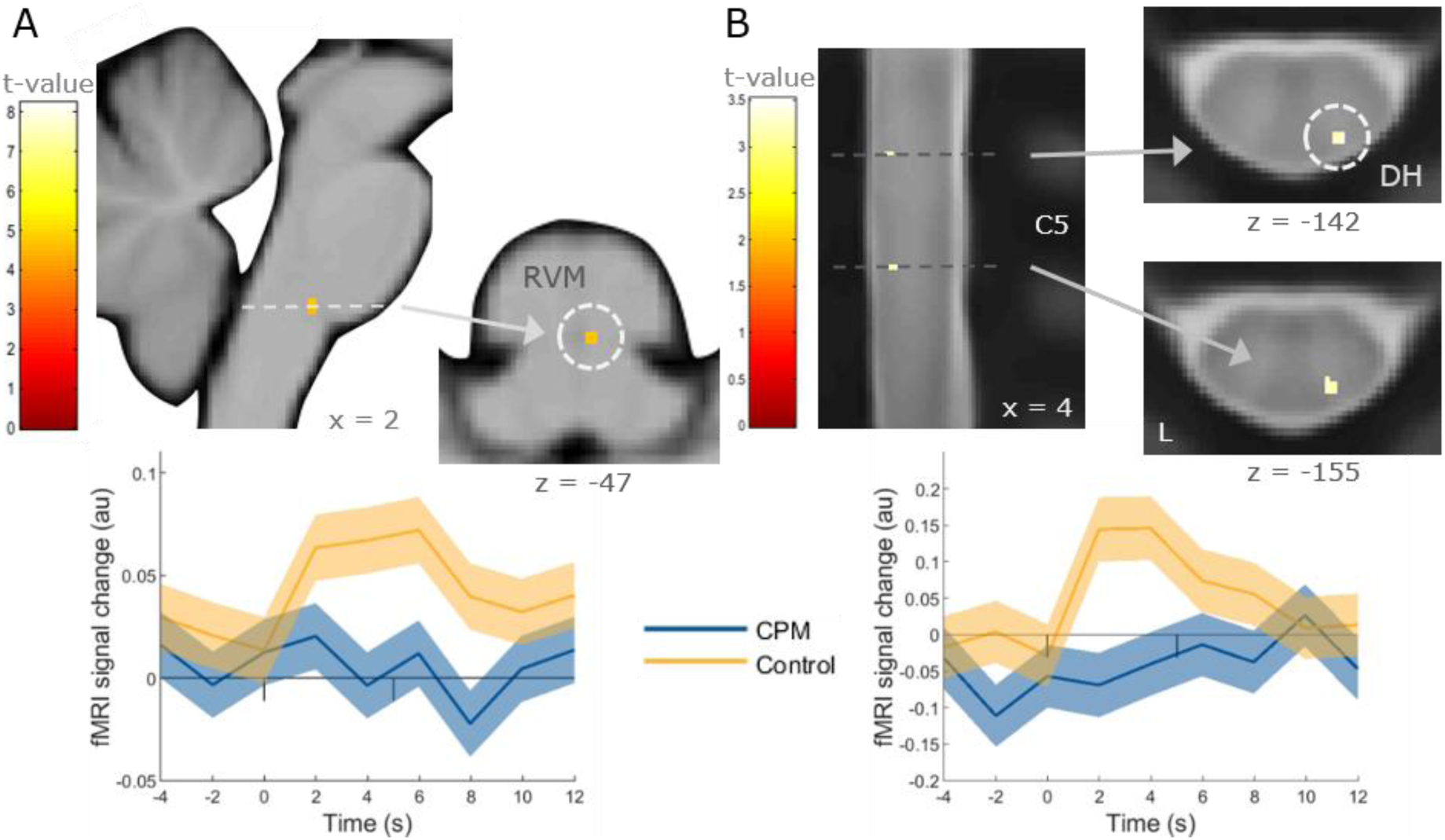
Pain-related regions in the brainstem and spinal cord showing CPM effects in a FIR model. (**A**) Time courses in the PAG and RVM show decreased activation in the CPM condition compared to the control condition. (**B**) Similarly, time courses in the right dorsal horn of the spinal cord show reduced activation in the CPM condition compared to the control condition. Both clusters are ipsilateral to stimulation site and at the height of the two vertebral discs spanning vertebral level C5. Pain-related region-of-interest masked statistical t-maps are overlaid on an average structural T1 image in MNI template space for A) and dorsal horn region-of-interest masked statistical t-maps are overlaid on a spinal mean EPI image inPAM50 template space for B). The visualization threshold is set to p_SVC_ < 0.05. Dashed circles indicate the cluster from which time courses of peak voxel parameter estimates are plotted. Shaded areas represent standard error of the mean. Ticks on the zero line indicate stimulus duration. Please note that the BOLD signal at a given time point reflects the averaged BOLD signal within one TR starting at that time point. RVM: rostral ventromedial medulla, DH: dorsal horn.

In a next step we tested whether areas of the descending pain modulatory pathway show the opposite activity pattern, namely increased activity in the CPM condition compared to the control condition. This analysis revealed two clusters in the vmPFC (xyz_MNI_: 6/45/-14, t_760_ = 4.71, p_SVC_ = 0.005; xyz_MNI_: − 10/54/-2, t_760_ = 5.15, p_SVC_ = 0.001) as well as a region in the vicinity of the PAG (xyz_MNI_: −9/-28/-9, t_760_ = 4.88, p_SVC_ = 0.002; Fig. 5), which showed increased activity in the CPM condition compared to the control condition.

**Figure 5.**
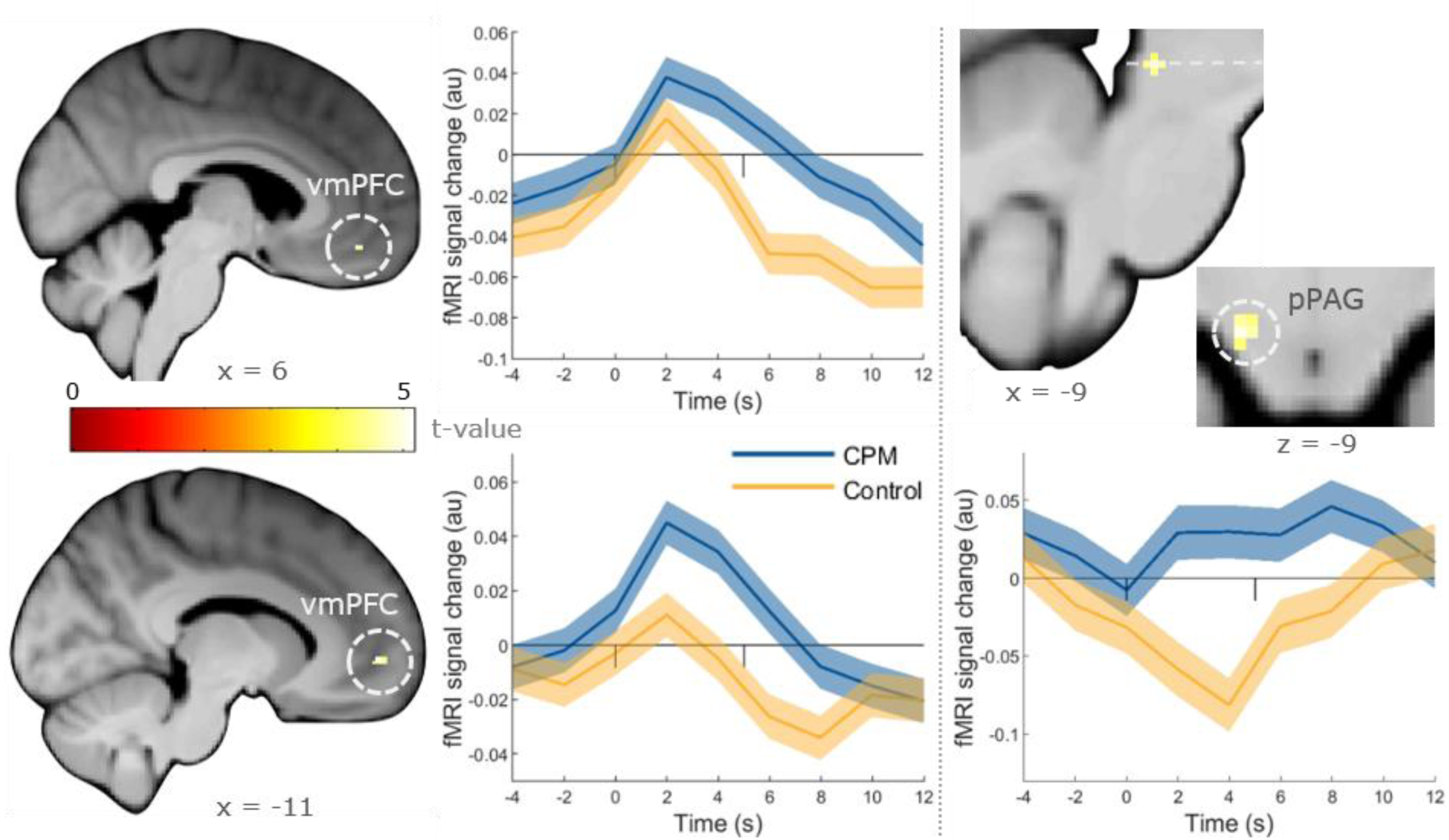
Regions of the descending pain modulatory system showing inverse CPM effects in a FIR model. Time courses in the vmPFC and brainstem show higher activation in the CPM condition compared to the control condition. Pain-modulatory region-of-interest masked statistical t-maps are overlaid on an average structural T1 image in MNI template space and the visualization threshold is set to p_SVC_ < 0.05. Dashed circles indicate the cluster from which time courses of peak voxel parameter estimates are plotted. Shaded areas represent standard error of the mean. Ticks on the zero line indicate stimulus duration. Please note that the BOLD signal at a given time point reflects the averaged BOLD signal within one TR starting at that time point. vmPFC: ventromedial prefrontal cortex, pPAG: peri periaqueductal gray.

Some regions which were either not included in our masks or which responded in a way that was contrary to our hypotheses showed further differences between CPM and control conditions. As we consider these results worth mentioning, we here report them as exploratory results. First, a region in the vmPFC/orbitofrontal cortex (xyz_MNI_: 3/44/-24, t_760_ = 6.05; Fig. S4A) showed reduced activity during the CPM condition mirroring responses in pain-related brain regions which was contrary to our hypothesis. Second, the primary somatosensory cortex which was not included in our pain modulation mask showed increased activation in the CPM condition. Strikingly, responses along the primary somatosensory cortex differed in that one cluster located more superior within the postcentral gyrus showed activation in both conditions with stronger activation in the CPM condition (xyz_MNI_: −29/-29/66, t_760_ = 5.96) while a second cluster located more inferior within the postcentral gyrus showed activation in the CPM condition but deactivation in the control condition (xyz_MNI_: −62/-6/29, t_760_ = 6.19; Fig. S4B).

### Representation of temporal CPM effects in the brain and spinal cord

In the next step, we investigated whether we could identify regions that show the temporal pattern of our behavioural CPM effect, namely an increase over the course of the experiment. Therefore, we tested which regions show an interaction of CPM versus control condition and time. This analysis revealed a cluster in the right dorsal horn of the spinal cord (xyz_MNI_: 5/-49/-153, t_39_ = 3.41, p_SVC_ = 0.046; Fig 6A). This cluster was located in the middle of vertebra level C5 along the z-axis. We also investigated whether regions along the descending pain modulatory pathway show the opposite effect, namely an increase in activity over the course of the experiment during the control condition. This analysis yielded a significant cluster in the vmPFC (xyz_MNI_: 3/33/-16, t_40_ = 4.55, p_SVC_ = 0.048; Fig 6B).

**Figure 6.**
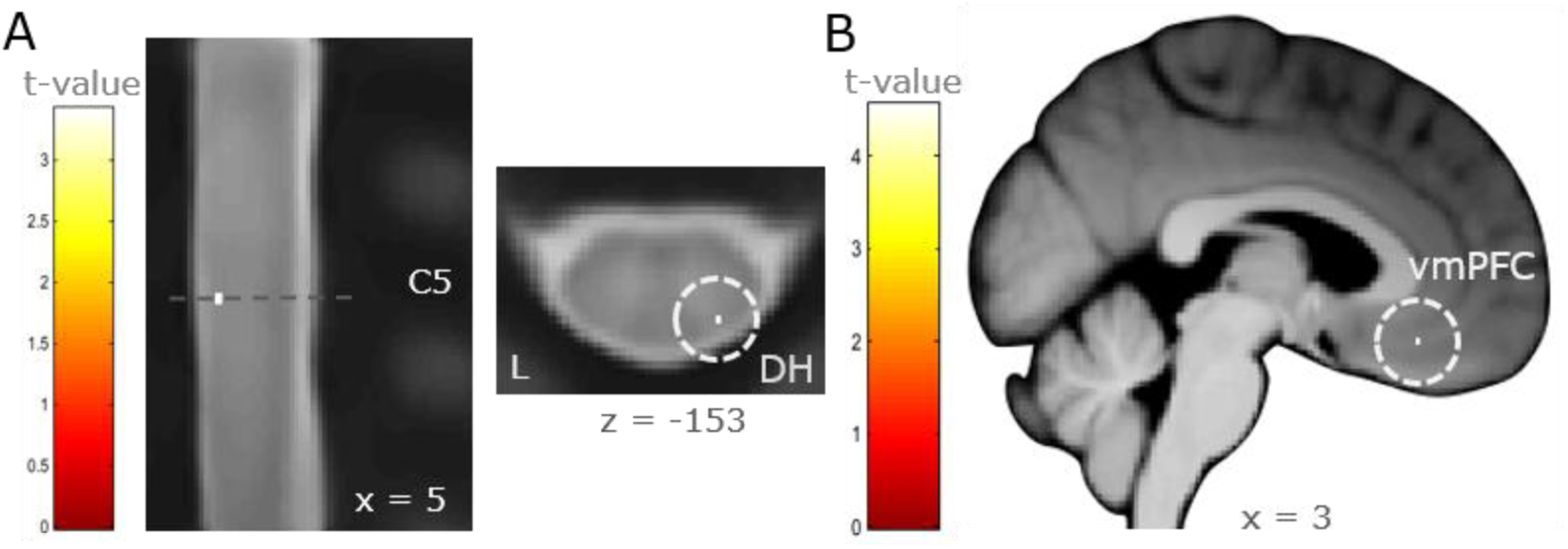
Temporal CPM effects. (**A**) The right dorsal horn in the spinal cord shows increasing activation over time in the control condition and decreasing activity over time in the CPM condition. (**B**) Along the descending pain modulatory pathway, a region in the vmPFC shows increasing activity over time in the CPM condition and decreasing activity over time in the control condition. Dorsal horn region-of-interest masked t-maps are overlaid on a spinal mean EPI image in PAM50 template space for A) and pain-modulatory region-of-interest masked statistical t-maps are overlaid on an average structural T1 image in MNI template space for B). The visualization threshold is set to p_SVC_ < 0.05.

### Neurological Pain Signature responses

In a next step we investigated whether CPM effects are reflected in brain wide pain-responsive regions thus combining pain-related and pain modulatory regions. To achieve this, we applied the neurological pain signature (NPS) (Wager et al., 2013), which is a multivariate fMRI-based marker for pain perception that assigns pain-responsive regions either positive (activated during pain) or negative (deactivated during pain) weights. Observing a distributed BOLD decrease across pain-responsive regions together with spinal attenuation would further indicate that CPM modulates bottom-up pain signalling. Again, this analysis was performed on the actual time courses of fMRI responses, showing that the NPS scores were significantly lower over time during the CPM condition compared to the control condition (mean±SE: 0.019±0.008, F[1,816] = 5.33, p = 0.021; Fig. 7A). Interestingly, responses were similar after stimulus onset with BOLD increases in both conditions that only differentiated between both conditions in the later part of the response, a pattern that was mainly driven by voxels with a positive weight within the NPS (Fig. 7B). Together with the finding that several pain-related brain regions are attenuated in the CPM condition, this temporal NPS pattern suggests that other pain-related brain regions show an increased BOLD response in the CPM condition. This observation motivated us to test whether pain-responsive brain regions represent both modulated and unmodulated pain signals. Therefore, we performed an equivalence test to identify voxels which show no meaningful difference between CPM and control condition. This analysis resulted in a binarized mask of voxels being statistically equivalent between conditions. These equivalent voxels were mostly localized contralateral to stimulation in the left postcentral gyrus (xyz_MNI_ = −26/-38/65), parietal operculum (xyz_MNI_ = −57/-30/21), central operculum (xyz_MNI_ = −41/-21/18) and anterior insula (xyz_MNI_ = −39/-6/11). Coordinates represent voxels from which time courses are plotted in Fig. 7C. Equivalent voxels were also located in the MCC (xyz_MNI_ = 8/20/35) and thalamus (xyz_MNI_ = −14/-17/9), although time courses were less conclusive (Fig. S5A). Equivalent ipsilateral responses were mostly restricted to the parietal operculum (xyz_MNI_ = 56/-29; Fig. S5B). Similarly, clusters in the vmPFC (xyz_MNI_ = −2/32/-8) and pgACC (xyz_MNI_ = 6/48/2) showed no meaningful BOLD difference between CPM and control condition (Fig. S5C), suggesting that modulated and unmodulated pain signals coexist within higher cortical pain-responsive brain regions.

**Figure 7.**
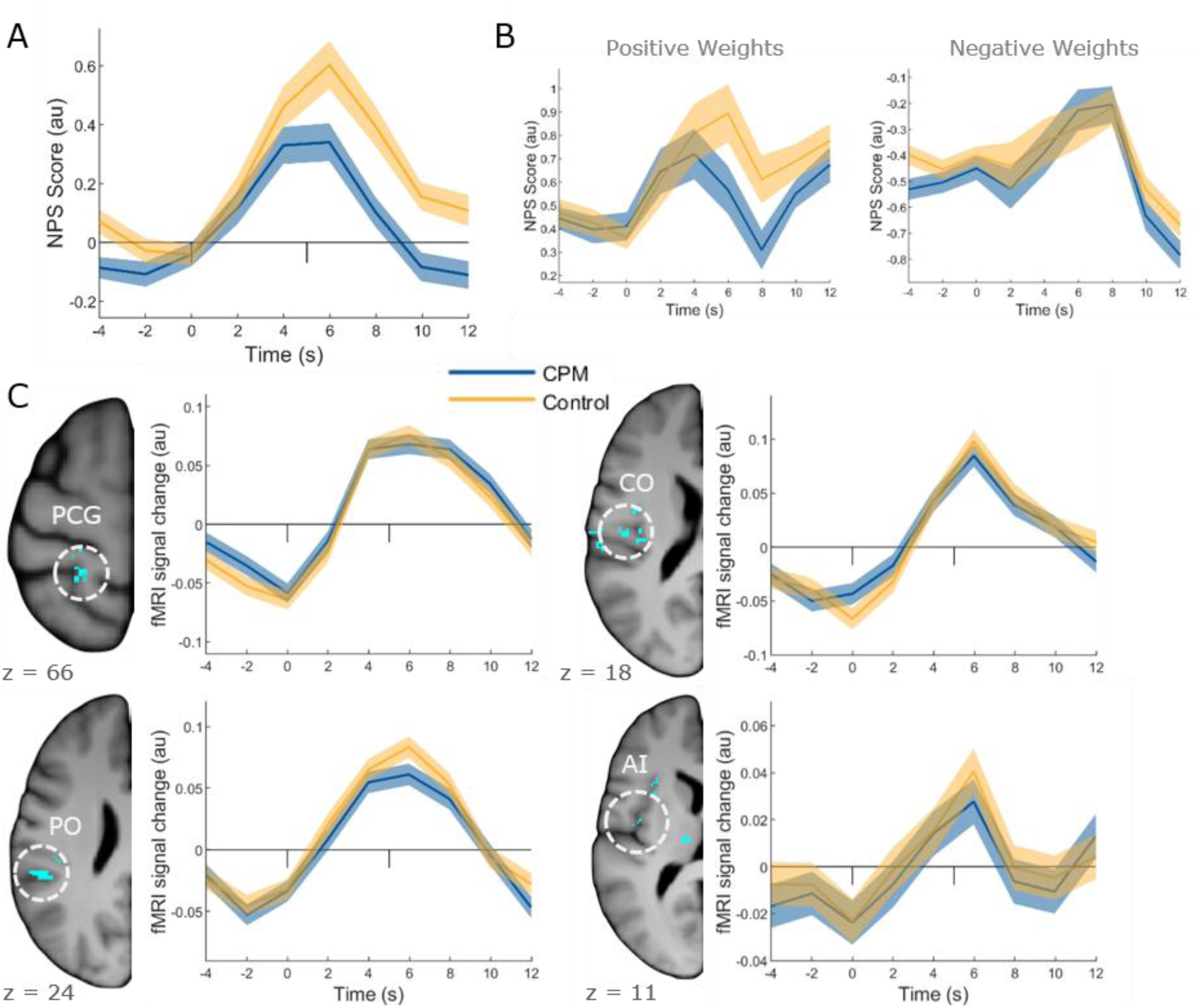
Neurological Pain Signature responses and equivalence between CPM and control. (**A**) NPS scores over time, derived from FIR analyses, show the most pronounced difference between CPM and control condition in the second half of the response. (**B**) NPS scores divided into positive and negative weights show that the overall NPS difference arises from brain regions contributing positive weights to the NPS. (**C**) An equivalence test shows no difference between CPM and control condition in some clusters within pain-related brain regions. Pain-related region-of-interest masked binary equivalence maps overlaid on an average structural T1 image in MNI template space. Please note that the BOLD signal at a given time point reflects the averaged BOLD signal within one TR starting at that time point. PCG: postcentral gyrus, PO: parietal operculum, CO: central operculum, AI: anterior insula.

### Coupling changes along the descending pain modulatory pathway

Last, we wanted to investigate whether the coupling between spinal cord and brain was modulated in the CPM condition. Therefore, we conducted a psychophysiological interaction (PPI) analyses with seed regions in the spinal cord, brainstem and vmPFC. The spinal cluster showing the CPM-related decreasing activity over the course of the experiment showed decreased coupling with the RVM (xyz_MNI_: 6/-38/-48, t_37_ = 5.21, p_SVC_ = 0.014) and two regions in the vmPFC (xyz_MNI_: −9/32/-10, t_37_ = 5.29, p_SVC_ = 0.012; xyz_MNI_: −2/40/-15, t_37_ = 5.41, p_SVC_ = 0.009; Fig. 8). In addition, the vmPFC showed decreased coupling in the CPM condition with another vmPFC region (xyz_MNI_: 6/33/-6, t_39_ = 6.38, p_SVC_ = 0.001) and with a brainstem region (not included in our mask) likely to be the subnucleus reticularis dorsalis (xyz_MNI_: −5/-41/-57, t_39_ = 4.36, Fig. S6A), which has been implicated in CPM (Bouhassira et al., 1992; Villanueva et al., 1996; Youssef et al., 2016b, 2016a). Conversely, PPI analyses originating in either RVM or PAG and targeting the spinal cord dorsal horn showed the following pattern: the RVM exhibited significantly decreased coupling with the spinal cord dorsal horn (xyz_MNI_: 3/-48/-148, t_38_ = 3.74, p_SVC_ = 0.022; Fig. S6B) while the PAG showed increased coupling with the spinal cord dorsal horn albeit not significant (xyz_MNI_: 3/-47/-155, t_38_ = 3.02, p_SVC_ = 0.104). Coupling strength did not correlate with individual CPM-induced analgesia across participants.

**Figure 8.**
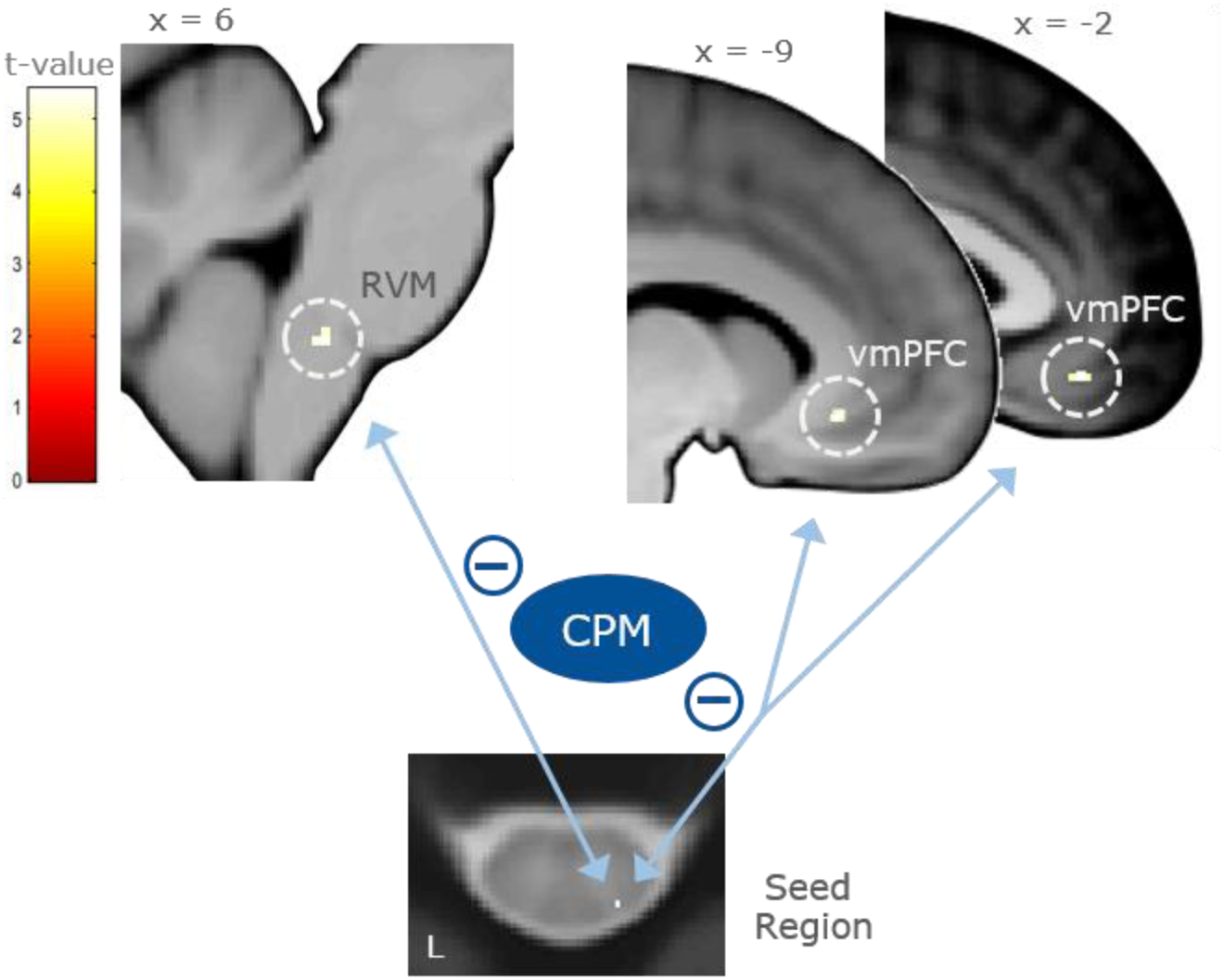
CPM-related coupling between the spinal cord and regions of the descending pain modulatory pathway. Coupling strength between the right dorsal horn, RVM, and vmPFC is decreased in the CPM condition compared to the control condition. Pain-modulatory region-of-interest masked statistical t-maps are overlaid on an average structural T1 image in MNI space and the visualization threshold is set to p_SVC_ < 0.05. RVM: rostral ventromedial medulla, vmPFC: ventromedial prefrontal cortex.

## Discussion

Our study provides a comprehensive investigation into the neural mechanisms underlying conditioned pain modulation by simultaneously imaging brain, brainstem, and spinal cord activity in humans. On the behavioural level, receiving painful phasic pressure stimuli with concurrent painful tonic pressure stimuli resulted in conditioned pain modulation over the course of the experiment compared to non-painful tonic control pressure stimuli. On the neural level, many pain-related brain regions such as the parietal and frontal operculum, anterior and posterior insula, mid-cingulate cortex, thalamus and RVM showed decreased activation in the CPM condition compared to control. Moreover, the right dorsal horn ipsilateral to the phasic stimulation site at spinal segment C6 showed reduced activity during the CPM condition. Contrary to pain-related regions, regions of the descending pain modulatory pathway such as the vmPFC and a brainstem region in the vicinity of the PAG showed increased activation during the CPM condition compared to control. The temporal CPM effect evolving over the course of the experiment was similarly mirrored in the right dorsal horn of spinal segment C6 while the opposite effect was represented in the vmPFC. Neural coupling along the descending pain modulatory pathway was further modulated under CPM as coupling between the spinal cord, RVM and prefrontal cortex was decreased compared to control.

### Behavioural effects

In this study, we did not find overall decreased pain ratings in the CPM condition compared to the control condition. One explanation is that CPM has been shown to depend on the degree of painfulness for both phasic and tonic stimuli (Chaudhry et al., 2025; Nir et al., 2011) and thus, it is possible that for some participants, the tonic and/or phasic stimuli were not painful enough to induce CPM. This is supported by the observation that the tonic rating explained some variance in the observed CPM effect. An alternative interpretation, namely carry-over effects between CPM and control blocks can be ruled out since there was no difference in pain ratings between participants that received control first and CPM second and participants that received CPM first and control second (see supplementary text). This suggests that CPM effects did not last longer than tonic stimulation and participants did not experience CPM in the control condition. Importantly, we added the control condition precisely to account for unspecific effects such as attention, whereas most CPM studies have compared phasic stimuli together with tonic stimulation to phasic stimuli alone. Given that attention to the conditioning stimulus has been shown to modulate CPM-related analgesia (Hoegh et al., 2019; Ladouceur et al., 2018, 2012) some part of the observed analgesia in previous CPM studies might be accounted for by these effects. Moreover, it is possible that heterotopic stimulation of different body parts is more potent to induce CPM effects than heterotopic stimulation of the same body part (Cummins et al., 2020; Hau et al., 2025; Oono et al., 2013). Many fMRI studies reporting robust CPM effects have used heterotopic stimulation with two different pain modalities such as a cold water bath as the conditioning stimulus and heat/pressure pain for the test stimulus (Bogdanov et al., 2015; Kisler et al., 2018; Nahman-Averbuch et al., 2014; Piché et al., 2009; Sprenger et al., 2011). However, another study using heterotopic pressure stimulation of the same body part similarly found a CPM effect over time (Harper et al., 2018), suggesting that stimulating the same body part produces more subtle CPM-related analgesia that accumulates over time.

### CPM-induced BOLD changes in pain-related brain regions and spinal cord

We found reduced activity under CPM in many pain-related brain regions such as the parietal operculum (SII), posterior and anterior insula, frontal operculum, thalamus, mid-cingulate cortex and RVM as well as in the spinal cord dorsal horn ipsilateral to stimulation site. These findings are in line with other fMRI studies that have similarly reported reduced activity across pain-related brain regions (Kisler et al., 2018; Piché et al., 2009; Song et al., 2006; Sprenger et al., 2011). In combination with reduced activity in the spinal cord, these findings hint to fact that spinal activity in the dorsal horn is inhibited under CPM and that this modulated response is then transmitted to brain regions involved in pain processing. This interpretation is also in line with theories about pain modulation which assume that modulated spinal signals are transmitted largely via the ascending spinothalamic tract to the thalamus and from there to cortical pain-related regions (Fields, 2004). Compared to other pain modulation conditions such as attentional or placebo analgesia, the widespread BOLD reductions across pain-related brain regions are noteworthy. Some studies report BOLD reductions in some of the pain-related regions for attentional (Bantick et al., 2002; Sprenger et al., 2012; Valet et al., 2004) and placebo analgesia (Eippert et al., 2009b; Kessner et al., 2013; Schenk et al., 2014; Wager et al., 2004; Wrobel et al., 2014; Zeidan et al., 2015). However, meta-analyses across placebo analgesia studies have shown that placebo analgesia does not systematically reduce activation across pain-related brain regions and that the anterior insula is the only region that most consistently shows reduced activity (Zunhammer et al., 2021, 2018). One conclusion might be that CPM involves a different descending pain modulatory mechanism compared to placebo or attentional analgesia which differentially affects activity across many pain-related brain regions. Another possibility is that pain modulation is more subtle than pain processing and shows a temporally different response pattern than stimulus-related responses. Although the motivation to focus on stimulus onset responses in this study was based on pressure pain results from a previous study, it is possible that our chosen time frame coincided with the temporal dynamics of CPM-related pain modulation. For example, some studies in the field of placebo research have found more pronounced placebo modulation in pain-related brain regions in the later phase of painful stimulation (Eippert et al., 2009a; Wager et al., 2004; Wrobel et al., 2014). This is in line with our observed temporal NPS responses that only show a distinction between CPM and control condition in the later part of the response. However, a systematic investigation of pain modulation responses across time is lacking so far.

Regarding laterality of phasic activity, we expected CPM effects to preferentially appear in the right dorsal horn ipsilateral to simulation site and in the left hemisphere contralateral to stimulation site.

While the right dorsal horn and left posterior insula showed this expected CPM effect, several regions such as the frontal operculum and anterior insula showed stronger CPM effects in the right hemisphere ipsilateral to the stimulation site which is similarly reported in other studies (Nahman-Averbuch et al., 2014; Piché et al., 2009; Sprenger et al., 2011). This might be explained by cross-hemispheric communication as pain often induces bilateral activity (Tracey, 2008), hence the right hemisphere may have received nociceptive input from the tonic stimulus together with cross-hemispheric input from the phasic stimulus.

### CPM-related temporal patterns

In this study, we focused on time course analyses since a previous study comparing heat and pressure pain indicated that cuff pressure-induced BOLD changes show a strong stimulus onset response without a sustained response throughout stimulus duration (Nold et al., 2025b). First evidence for such a pressure pain-related pattern comes from an earlier study (Loggia et al., 2012) which is in stark contrast to heat pain which induces BOLD responses that mirror stimulus profiles (Horing et al., 2019; Tinnermann et al., 2022). In the study by Nold and colleagues, stimuli lasted about 15 seconds while the BOLD response was only present for approx. 8 seconds. Results in this study confirm this initial observation as time courses in the parietal operculum, posterior insula, and RVM showed a similar response pattern. This suggests that traditional HRF-based GLM analyses are not well suited to pressure pain studies.

To our knowledge, this is the first study to show pain-related time courses in the spinal cord. One reason why we were able to detect stimulus-related time courses in the spinal cord might be due to the pressure stimulation device that was used in this study. Heat pain studies often stimulate a very small skin area while the pressure cuff stimulates a larger area. Stimulation of a larger area should recruit more sensory and nociceptive neurons and thus might lead to a more robust activation in the spinal cord that is more sensitive to time course analyses. Another interesting aspect about the spinal cord time course is the unusually fast and steep increase after stimulus onset. While a comparison in HRFs between the brain and spinal cord in non-human primates did not show a difference between spinal cord and brain hemodynamic responses for heat pain (Yang et al., 2015), the early response onset in this study raises the question whether a traditional HRF is an appropriate assumption to model spinal cord responses in pressure pain. More studies investigating time courses in the spinal cord are necessary to see whether this pattern is specific for pressure pain or generalizes to other pain modalities.

### Descending pain modulatory system responses

In line with our hypothesis, we found that regions of the descending pain modulatory pathway such as the vmPFC showed the opposite activity pattern, i.e. increased activity in the CPM condition compared to the control condition. A similar response was observed in the vicinity of the PAG. Moreover, the vmPFC displayed the opposite temporal CPM effect. Increased prefrontal activity under CPM has been reported in other fMRI studies investigating CPM effects (Kisler et al., 2018; Nahman-Averbuch et al., 2014; Sprenger et al., 2011) and corroborates findings from studies investigating pain modulation processes such as placebo- and attention-based analgesia that similarly report increased vmPFC activity during analgesic modulation (Atlas et al., 2012; Bantick et al., 2002; Eippert et al., 2009a; Geuter et al., 2013; Kong et al., 2009; Koyama et al., 2005; Petrovic et al., 2002). This is particularly interesting as CPM effects do not depend on cognitive influences or learning as compared to placebo effects and still involve the vmPFC in a similar fashion. Responses in the brainstem are more complex as in this study, the RVM showed a CPM effect while the vicinity of the PAG showed an inverted CPM effect. Some previous studies have found that during pain modulation the PAG can mirror behavioural responses (Atlas et al., 2012; Crawford et al., 2025, 2021; Tinnermann et al., 2022), while others have found inverted PAG responses (Eippert et al., 2009a; Petrovic et al., 2002; Scott et al., 2008; Tracey et al., 2002). Similarly conflicting results are reported for the RVM with decreased activity (Crawford et al., 2025, 2021; Sprenger et al., 2011) as well as increased activity (Brooks et al., 2017; Eippert et al., 2009a) during pain modulation. These results suggest that modulatory brainstem nuclei can respond in both directions and more research is needed to investigate whether these different responses are spatially localized and in which conditions these nuclei respond one way or the other.

### CPM-related responses in the primary somatosensory cortex

That primary somatosensory cortex activity was higher in the CPM condition compared to control is particularly interesting. Other studies have similarly reported increased SI activity during CPM (Kisler et al., 2018; Nahman-Averbuch et al., 2014) which could result from the larger amount of sensory and nociceptive fibres being stimulated during the CPM condition compared to the phasic stimulus alone. The second more inferior cluster in SI showed a deactivation in the control condition and an activation in the CPM condition therefore mirroring prefrontal and PAG responses. This implies that some subregion within the primary somatosensory cortex responds with deactivations to nociceptive stimulation.

### CPM-related Neurological Pain Signature responses and equivalence

Previous research has indicated that pain modulation such as placebo analgesia does not lead to modulation of neural responses in pain-related brain regions since NPS scores during placebo conditions were not significantly decreased across studies (Zunhammer et al., 2018). Here, we provide first evidence that CPM is associated with reduced NPS scores indicating decreased BOLD responses across several pain-related brain regions. Together with inhibited spinal activity under CPM and decreased coupling along the descending pain modulatory pathway, these findings could be interpreted as evidence for descending pain modulation of spinal activity and subsequent bottom-up signalling of modulated spinal responses to different pain-related brain regions. However, the temporal NPS score showed a substantial increase during the CPM condition thus suggesting that a considerable amount of voxels in pain-related brain regions still responds to the phasic stimulus in the CPM condition. Therefore, we performed an equivalence test to determine which voxels in pain-responsive regions do not show a difference between CPM and control condition. This analysis revealed some clusters in cortical pain-related regions such as the primary and secondary somatosensory cortex and the operculum that responded in a similar fashion to phasic stimulation irrespective of condition. This finding indicates that regions which are targeted first by ascending pathways, represent both the unmodulated as well as the modulated pain signal.

### CPM-related neural coupling across pain modulatory pathways

We found decreased coupling between the spinal cord dorsal horn and regions of the descending pain modulatory pathway, namely the vmPFC and RVM. Coupling changes along pain modulatory pathways have been reported in several other CPM studies. In task-based studies, CPM-related coupling between the prefrontal cortex, PAG and RVM was increased (Piché et al., 2009; Sprenger et al., 2011). Similarly, resting-state studies have reported coupling changes between the prefrontal cortex and the PAG (Harper et al., 2018; Li et al., 2025). Some of the studies have further reported associations between coupling strength change and individual CPM effect magnitudes (Li et al., 2025; Sprenger et al., 2011). These CPM-related coupling patterns are in line with other modulation studies such as attention and placebo-induced analgesia that report coupling changes between the prefrontal cortex and the PAG (Bingel et al., 2006; Eippert et al., 2009a; Petrovic et al., 2002; Valet et al., 2004; Wager et al., 2007). These findings indicate that CPM relies on the same descending mechanism as other pain modulation processes to ultimately influence spinal cord activity. It is however surprising that the coupling strength was decreased under CPM as we expected to find increased coupling between regions of the descending system. Taken together with our (non-significant) finding of increased coupling between PAG and spinal cord, our observation could imply that pain modulation relies on different coupling patterns along the descending pain modulatory system.

### Limitations

Experimental CPM paradigms for fMRI studies have some limitations regarding control of confounding factors. First, parallel application of phasic and tonic stimuli has the disadvantage that CPM-related BOLD responses can never be fully attributed to the phasic stimulus alone. However, a sequential application of tonic and phasic stimuli is not suitable as it is not known how long CPM effects last after tonic stimulus application (Imai et al., 2016; Nahman-Averbuch et al., 2013) and most behavioural studies using a sequential protocol assess CPM on pain thresholds directly after tonic stimulation and not on pain perception during repeated painful stimulation. Moreover, parallel stimulation is advantageous over sequential stimulation as it results in greater CPM effects (Hau et al., 2025; Pud et al., 2009; Sirucek et al., 2020) and is further recommended in order to induce modulatory mechanisms that might be similar to DNIC in animals (Piché, 2023). Second, an appropriate control for the tonic stimulus is difficult to establish. Many studies do not implement a control condition and compare the tonic plus the phasic stimulus to the phasic stimulus alone. In this study, we tried to implement a control condition to account for sensory (Hollins et al., 2014; Lundeberg et al., 1984) and attention effects (Hoegh et al., 2019; Ladouceur et al., 2018, 2012) during parallel stimulation, as both factors might otherwise contribute to the observed analgesic effect. Nevertheless, a long-lasting painful stimulus still involves larger sensory afferent drive and captures more attentional resources than a non-painful stimulus.

### Conclusions

In the present study, conditioned pain modulation was associated with widespread decreased BOLD responses in pain-related brain regions and the spinal cord dorsal horn. Furthermore, increased BOLD responses in the CPM condition were present in pain-modulatory brain regions such as the vmPFC and in the vicinity of the PAG. Coupling across the descending pain modulatory pathway was modulated under CPM with decreased coupling between the spinal cord dorsal horn, RVM and vmPFC. These results suggest that CPM involves descending modulation originating in the vmPFC to attenuate spinal dorsal horn activity and that this attenuated spinal signal is then conveyed to pain-related brain regions. The results of our cortico-spinal imaging study bridge the gap between existing human fMRI studies investigating CPM effects in the brain and rodent studies investigating brainstem-spinal circuits to characterize descending mechanisms during DNIC.

## Materials and Methods

### Participants

Fifty healthy and right-handed participants took part in the study. Inclusion criteria were a body-mass-index (BMI) between 18 and 30, no contraindications to MRI, no acute or chronic pain, no acute or chronic somatic or psychiatric illness, no recent or chronic injury in the arms, no current muscle soreness or large bruising in the arms, and no regular intake of medication (except for contraceptive pill, allergy medication, or medication for hyper- or hypothyroidism). Participants were asked to refrain from intensive physical exercise on the day of the experiment and to not take pain killers in the 24 hours preceding the experiment. The study was approved by the Ethics Committee of the Medical Council of Hamburg (2020-10144-BO-ff) and conducted according to the Declaration of Helsinki. Written informed consent was be obtained from all participants prior to study participation, and after the study, participants were compensated with 45-55€ depending on study duration (usually 2 hours). Eight participants had to be excluded. One participant was excluded because of very low pain ratings for the tonic stimulus, four participants aborted the experiment, one participant fell asleep, and two participants were excluded because of technical issues. The final sample consisted of 42 participants (21 women, 21 men, mean age 26.7 ± 5.3 years, mean BMI 23.7 ± 3.2 kg/m^2^).

### Pressure stimuli

Tonic and phasic stimuli were implemented through parallel and heterotopic stimulation of the upper arms with the same pain modality. Parallel stimulation is applied in many CPM-related fMRI studies, has been proposed as a requirement to reflect a DNIC-like mechanism in humans and has been shown to induce stronger CPM-related analgesia as sequential stimulation (Hau et al., 2025; Pud et al., 2009; Sirucek et al., 2020). Although heterotopic stimulation of different body parts is often recommended in the literature (Piché et al., 2009; Sirucek et al., 2023; Yarnitsky et al., 2015), we implemented heterotopic stimulation of the same body part as our cortico-spinal imaging approach does not allow measuring large field of views in the spinal cord which would be required for stimulating different parts (e.g. arm and leg). In comparison to previous studies that compared tonic plus phasic stimuli to phasic stimuli alone, we furthermore implemented a control condition (non-painful pressure stimuli) to account for unspecific sensory stimulation and attention effects.

Pressure stimuli were delivered to the right and left upper arm with a computerized cuff algometry device (CPAR, Nocitech, Denmark). The conditioning stimulus was a tonic pressure stimulus on the left arm and the test stimulus was a phasic pressure stimulus on the right arm (Fig. 1B). The tonic stimulus lasted for 200 s and was either a non-painful control stimulus or a painful conditioning stimulus. The tonic stimulus had a sinusoidal cyclic form with three repetitions of peaks and troughs (60 seconds per cycle) and a ramp up and ramp down phase in the beginning and end of 10 seconds. For the non-painful tonic control stimulus the sinusoidal form was realized through perceptible pressure fluctuations between 2 and 5 kPa. For the painful tonic stimulus, pressure levels were individually calibrated to fluctuate between 50 and 70 VAS. Phasic painful pressure stimuli lasted 5 seconds and were individually calibrated to 60 VAS. Nine phasic stimuli were distributed around predefined time points along one tonic stimulus. These time points were at 5, 20, 35 and 50 seconds of one sinusoidal cycle resulting in 12 possible time points throughout a tonic stimulus where phasic stimuli could appear. The 9 phasic stimuli were randomized to these 12 possible time points with the constraint that each cycle always included 3 phasic stimuli. Furthermore, the real onset of the phasic stimuli was jittered between 0 and 2 seconds around the predefined time points. The inter-stimulus interval between phasic stimuli was between 8 and 37 seconds.

### Pressure stimulus calibration

Calibration of the pressure stimuli took place while participants were lying in the MRI scanner. Tonic stimuli (left arm) and phasic stimuli (right arm) were calibrated separately but the procedure was identical for both stimulus types. First, participants received two low pressure stimuli (10 and 20 kPa) to familiarize them with pressure cuff stimuli. Next, participants received six stimuli around an average pain threshold (based on pilot studies) and rated them as either non-painful or painful. The pain threshold was defined as the lowest pressure that was rated as painful. The pain threshold estimation was realized for both stimulus types/arms before continuing with the other parts of the calibration and the order was counterbalanced between participants. For the following part of the calibration, participants were familiarized with the Visual Analog Scale which consisted of 101 rating steps from 0 to 100 where 0 was instructed to be non-painful, and 100 to be pain experienced as intolerable for an extended duration in the context of the experiment. Next, participants experienced four predefined stimulus intensities that were based on their pain threshold and three intensities that were inferred from their ratings of those four stimuli. A linear regression was fit to the intensities and their respective ratings and a variable number of stimuli was applied until the estimation of the linear regression was reliable enough to infer pressure levels that corresponded to 50 and 70 VAS for the tonic stimulus/left arm and 60 VAS for the phasic stimulus/right arm. The calibration order for stimulus type/arm was counter-balanced across participants.

### Experimental design

The experiment consisted of 4 experimental runs during which BOLD responses in the brain and spinal cord were acquired. One run consisted of two tonic stimuli that always belonged to the same condition (painful or non-painful). The two tonic stimuli were separated by an inter-stimulus-interval of 30 seconds. Nine phasic stimuli distributed across one tonic stimulus resulted in 18 phasic stimuli per run. Phasic stimuli in the innocuous condition are referred to as control stimuli whereas phasic stimuli in the noxious condition are referred to as CPM stimuli. In total, two runs per condition were acquired resulting in 4 tonic stimuli and 36 phasic stimuli per condition. Conditions were counterbalanced across participants. Throughout the experiment, there was a white fixation cross on a grey background except during rating periods. A VAS rating scale appeared after a variable delay between 0 and 1 s after phasic stimulus offset and lasted for 5 seconds. Right before the first fMRI run and directly after the last fMRI run, participants received 80 seconds of the tonic stimulus (one cycle plus ramp and ramp down) and continuously rated its intensity of the stimulus. We chose 80-second stimuli for rating because pilot testing showed that participants reached a rating plateau after around 60 seconds. These ratings show that the tonic stimulus was perceived as painful throughout the experiment as pre and post stimuli were rated on average as medium painful (Fig. S1A). However, the painfulness of the tonic stimulus increased over time as during the first 50 seconds of stimulus duration, participants rated the post stimulus as more painful than the pre stimulus. Verbal reports after the experiment indicated that participants perceived numbness and “pins-and-needles” sensations in the arm which might be caused by ischemic effects as the pressure cuff obstructs blood flow to the rest of the arm (Graven-Nielsen et al., 2015) and which could explain the observed sensitization.

### Physiological measures

Stimulus presentation, pressure device triggering and response logging were realized with MatlabR2020a and Psychophysics Toolbox 3.0.17 (http://psychtoolbox.org). Physiological data were recorded at 1000 Hz during scanning with the Expression system (In Vivo, Gainesville, US) and a respiration belt around the chest and a pulse oximeter at the foot. Physiological signals were further digitized together with scanner pulses by a CED1401 system and spike2 software (Cambridge Electronic Design). For post-processing of data, respiration and pulse signals were downsampled to 100 Hz, detrended and smoothed (pulse with 5 and respiration with 50 samples FWHM Gaussian kernel).

### MRI data acquisition

In this study, we measured BOLD responses simultaneously in the brain and spinal cord, which has successfully been employed in previous studies (Landelle et al., 2024; Oliva et al., 2022, 2021; Pfyffer et al., 2025, 2024; Sprenger et al., 2015; Tinnermann et al., 2022, 2017; Vahdat et al., 2020, 2015). This approach allows investigating neural responses across the entire central nervous system including the spinal cord to characterize cortico-spinal interactions during pain perception and pain modulation. Here, we applied the imaging approach proposed by Finsterbusch and colleagues (Finsterbusch et al., 2013) that allows defining two (brain and spinal cord) subvolumes with different geometric resolutions and timings. Prior to scanning, we determined optimal shim parameters separately for brain and spinal subvolumes using a standard field map acquisition and a dedicated shim algorithm as described recently (Chu et al., 2023) that were dynamically updated during EPI measurements (Blamire et al., 1996; Morrell and Spielman, 1997). During the shimming procedure, which further included measurements of T1 and T2-weighted anatomical images, participants completed the pressure stimulus calibration.

All MRI acquisitions were performed on a 3 Tesla whole-body MR system (Magnetom PrismaFit, Siemens Healthineers, Erlangen, Germany) using a standard 64-channel head-neck coil. The isocenter of the magnet was set to the chin of participants. During the experiment, T *-weighted echo-planar imaging (EPI, flip angle 70°, TR 1991 ms, descending slice acquisition order) covering 60 slices without gap in the brain (FOV 220 x 220 mm^2^, voxel size 2.0 x 2.0 x 2.0 mm^3^, TE 24 ms, 56.0 ms per shot) and 12 slices without gap in the spinal cord (FOV 132 x 132 mm^2^, voxel size 1.2 x 1.2 x 5.0 mm^3^, TE 27 ms, 72.6 ms per shot) was implemented. Simultaneous multi-slice imaging (blipped-CAIPI) was applied for the brain slices with an acceleration factor of 3 (Chu and Finsterbusch, 2021). In the spinal cord, optimal z-shim parameters were determined for every slice (Finsterbusch et al., 2012), and a saturation pulse was applied to the throat and chin region minimizing ghosting and inflow artefacts related to pulsatile blood flow in the major cervical vessels. Adjustment volumes covered the full brain EPI volume and a 40 x 40 x 60 mm^3^ volume in the spinal cord, respectively. The brain field-of-view was aligned with the bottom-most slice including rostroventral medulla below the pons and the cerebellum in its entirety. Consequently, in some participants the top of the brain including primary somatosensory cortex was not fully included. For the spinal cord, the topmost slice was positioned to the upper edge of the spinal disc C3-C4.

Anatomical images included a high-resolution structural T1-weighted scan covering the entire head and the neck up to the lower part of the second or upper part of the third thoracic vertebra, depending on participant height (MPRAGE; TR 2300 ms, TE 3.41 ms, inversion time 1100 ms, 1.0 x 1.0 x 1.0 mm^3^ voxel size, flip angle 9°, FOV 320 x 240 x 192 mm^3^, sagittal slice orientation), and a T2-weighted acquisition of the spinal cord including vertebrae C2-T2 (3D turbo spin echo, TR 1500 ms, TE 120 ms, 0.8 x 0.8 x 0.8 mm^3^ voxel size, refocusing flip angle 120°, FOV 256 x 256 x 51 mm, 64 sagittal slices).

### Behavioural data analyses

Behavioral data were analyzed using t tests and linear mixed effects models as implemented in MATLAB’s *fitlme* function (The MathWorks, Massachusetts, USA; version MatlabR2023a). A paired t test was calculated to assess differences between phasic pain ratings in the control and CPM condition. The time effect of CPM-induced analgesia was modelled using a linear mixed effects model with trial-wise phasic pain ratings as dependent variable, subject number as random effect and the condition and trial number (from 1 to 72, then mean-centered) as fixed effects.

The influence of the painfulness of the tonic stimulus on CPM was assessed using a linear mixed model with trial-wise pain ratings as dependent variable, subject number as random effect and the condition and mean-centered tonic rating as fixed effects. For the latter, we averaged the continuous rating of the tonic stimulus right after the experiment excluding ramp times of 10s in the beginning and end of the stimulus (Fig. S1B).

### Brain preprocessing

After acquisition, DICOM files were converted into NIfTI files, separately for the brain and the spinal cord sub-volumes. Brain and spinal cord subvolumes were separately preprocessed and analysed. Brain fMRI data were analysed with SPM12 (Wellcome Trust Centre for Neuroimaging, London, UK). First, the origin in brain images was relocated to the brain which helps normalization since the origin in combined imaging lies in the neck. Next, EPI images were realigned and the T1 image was segmented and normalized with DARTEL to the MNI152NLin2009cAsym template brain in Montreal Neurological Institute (MNI) standard space. The mean EPI was coregistered to the T1 using a non-linear algorithm. Next, the realignment of EPI images was repeated (on raw images) using a mask that excluded the eye region to reduce the bias of eye movement in motion correction. Last, EPI images were normalized using the DARTEL flow fields. Smoothing of images with a 6 mm FWHM kernel was done after first-level analyses with the resulting contrast images. One participant had to be excluded from imaging results due to artefacts in functional images.

### Spinal cord preprocessing

Spinal cord data were preprocessed with the Spinal Cord Toolbox version 6.0 (De Leener et al., 2017) and SPM24. The anatomical T2 image of the spinal cord was segmented with a deep learning model (Gros et al., 2019). Next, the vertebrae were automatically labelled and T2 images were registered to the PAM50 template space using the segmentation as well as the vertebrae labels (De Leener et al., 2018). EPI images within run were realigned using the slice-wise realignment method in SPM24. All subsequent preprocessing steps were realized in the Spinal Cord Toolbox. Mean EPI images for each run were registered to the mean EPI of the first run in a first step and to the mean EPI image of all runs in a second step similar to the realignment realized in SPM. Subsequently, the spinal cord of the resulting mean EPI was segmented using a newly developed algorithm for functional images (Banerjee et al., 2025) to help the registration of the mean EPI image to the T2 image. Mean EPI images in T2 space were normalized to the PAM50 template space using the T2-to-template-space warping field as initial registration step. Finally, EPI images were warped using a warping field that combined all registration steps from registration between runs to registration to T2 space and to PAM50 template space. After first-level analyses, contrast images were smoothed with a 1 mm FWHM kernel. Please note that in all registration steps, registration parameters were adapted for some participants to obtain better registration results. Registration results were inspected visually to determine a successful registration. To assess the quality of the spinal data, temporal signal-to-noise (SNR) ratios within the spinal cord were calculated per participant for every slice after motion correction (Fig. S7). For visualization purposes, the temporal SNR maps were warped to template space and averaged across all participants (Fig. S7). Another participant had to be excluded from all spinal analyses because the T2 normalization failed due to bad data quality.

### First-level statistical analyses

For main statistical analyses, “post-stimulus averaging” models were defined using a Finite Impulse Response function separately for brain and spinal cord. This type of model was chosen based on a previous study from our lab comparing heat and pressure pain (Nold et al., 2025b). Temporal response patterns for pressure pain showed an initial stimulus onset response that rapidly decayed. This response pattern differs from heat pain as it does not show a stable activation level throughout stimulus duration. Thus, convolving task regressors with a hemodynamic response function might include inaccurate assumptions. The bin width was set to the TR and ten bins were modelled for control and CPM conditions separately. The onset of regressors was set to two bins prior to phasic stimulus onset. Nuisance regressors for the brain included 24 motion regressors (derivatives of the initial 6 motion parameters from realignment, and squares of the original and derivative parameters), 18 RETROICOR regressors for cardiac and respiratory noise (4 orders of cardiac, 3 orders of respiratory and 1 order of interaction components) (Chang et al., 2009; Glover et al., 2000), and 24 principal component regressors to model noise arising from white matter (WM) and cerebrospinal fluid (CSF), respectively.

Nuisance regressors for the spinal cord included 6 motion regressors for spinal movement, 24 motion regressors from the brain volume, 18 RETOICOR regressors and WM and CSF regressors. For the CSF regressors, a variance image across all EPI images in one run was calculated that captured the CSF region quite precisely. This image was masked with the segmented spinal cord of the mean EPI to exclude all voxels within the spinal cord and finally, a principal component analysis was calculated within the resulting CSF mask. All components that explained 95% of the variance within one run were included as nuisance regressors. For white matter regressors, gray matter was segmented from individual mean EPI images and dilated by 2 mm. The resulting images were cut from the individual segmented spinal cord and the resulting white matter mask was used for a principal component analysis. Here, all components that explained 50% of the variance within one run were included as nuisance regressors. Moreover, we included censoring regressors for those volumes that exceeded a certain threshold with regard to deviation from mean voxel intensity which identifies volumes with excessive motion or imaging artefacts. All first-level analyses were filtered with a high-pass filter of 240 seconds. After model estimation, first-level contrast images were calculated for every bin and participant.

Additional conventional first-level models were defined that included task regressors for the experimental conditions ‘phasic CPM’, ‘phasic control’, ‘rating’, ‘tonic painful’ and ‘tonic control’. Phasic stimuli were modelled as a 5 s boxcar function and tonic stimuli as a 200 s boxcar function. The rating was separated into two regressors for brain analyses as the rating scale always appeared for 5 s. Hence, the rating duration was set to the reaction time and the remaining time was modelled as visual scale regressor. This regressor was not included into spinal analyses as visual input is not expected to have any influence on the spinal cord activity (for the corresponding results, see supplementary text and Fig. S8). To test temporal developments in CPM effects across the course of the experiment, an additional concatenated first-level model including one regressor for ‘phasic’ was created. Here, a parametric modulator combining the phasic trial number (1-72) and the condition (CPM or control) was added to the model representing the interaction between condition and time. Nuisance regressors were identical to the first first-level approach. Experimental regressors were convolved with the canonical haemodynamic response function. After model estimation, the first-level contrast images ‘phasic’, ‘phasic control’, ‘control>CPM‘ were defined for each participant (first model) and ‘time*condition modulator’ (second model).

### Second-level analyses

The resulting contrast images from each participant were then raised to random-effects group analyses using a flexible factorial design for FIR models and one-sa9mple t tests for HRF models. The flexible factorial design included factors for ‘subject’, ‘condition’ and ‘FIR bin’. Second-level contrasts of interest for FIR models accounted for differences between CPM and control conditions in the first 8 seconds after stimulus onset based on pressure-related time courses from our previous study (Nold et al., 2025b). To test for correlations between BOLD responses and behavioural CPM effects difference ratings between CPM and control conditions were included as covariate in second-level t tests.

### Neurological Pain Signature analyses and equivalence tests

To examine the compound activity across all voxels in pain-responsive pain regions, we applied the Neurological Pain Signature which is a brain-based biomarker for pain that assigns different weights to voxels based on their contribution to predict pain perception (Wager et al., 2013). To do so, we multiplied all 10 FIR contrast images for both conditions (CPM and control) with the NPS mask.

Differences were assessed using a linear mixed model with condition and FIR bin as predictors. We additionally multiplied FIR bins with either positive or negative voxel weights of the NPS to assess the contribution of pain-responsive activated versus deactivated voxels. Moreover, we multiplied contrast images from a HRF model for control and CPM conditions with the NPS and the Stimulus Intensity Independent Pain Signature (Woo et al., 2017) mask and assessed significance with paired t-tests (for corresponding results, see supplementary text and Fig. S9). The results from FIR-NPS analysis motivated us to identify brain regions that show no difference between CPM and control condition. Therefore, we performed an equivalence test based on a method that derives the effect size g as well as the upper and lower bound of the confidence interval for every voxel from a t-map (Gerchen et al., 2021). Here, we used a second-level paired t test between phasic control and phasic CPM contrast images to calculate effect sizes and confidence intervals. We defined our equivalence criterion as a confidence interval crossing 0 and not overlapping with an equivalence bound of ±0.4 which resulted in a binarized mask of voxels showing significant equivalence between CPM and control condition. As this analysis reveals many noisy voxels, we restricted our criterion to voxels located within our pain mask (see section ‘Multiple comparisons correction’) that show pain-related activity defined through a thresholded second-level contrast for a main effect of phasic control stimulation. The resulting voxels display effect sizes (Hedges g) between −0.036 and 0.04 with a mean t-value of 0.009±0.09 ranging between −0.164 and 0.166. For pain-modulatory regions, we masked the equivalence criterion map with our descending pain mask. Time courses based on the FIR model were plotted for some of the voxels to visually show that responses in these voxels were similar between conditions.

### Connectivity analyses

To characterize the functional integration between regions of the modulatory pain system we implemented psycho-physiological interaction (PPI) models with seed regions in the spinal cord (Tinnermann et al., 2022, 2017). These seed regions were defined based on results from previously described second-level analyses. Accordingly, we extracted individual time courses from three regions-of-interest (ROI) in the spinal cord that were derived from two spinal clusters that showed reduced activity under CPM compared to control and from one cluster that mirrored behavioural effects with a decrease of pain ratings in the CPM condition over time. The centre of these ROIs was set to the peak voxels of the respective clusters and the radius was set to 2 mm. Time courses within these ROI were extracted from voxels that exceeded the statistical threshold of 0.5 in the effects-of-interest contrast. Additional seeds in the RVM, PAG and vmPFC were defined based on second-level FIR results. Seeds in the brainstem had a radius of 3 mm and 4 mm in the vmPFC. The same threshold for the effects-of-interest contrast was used as for the spinal seeds.

Since the estimation of functional connectivity can be influenced by physiological noise (Birn, 2012) and movement (Van Dijk et al., 2012), the extracted time courses were adjusted for variance explained by nuisance regressors such as the mean, RETROICOR and movement. The PPI models included three regressors, one for the psychological variable (control>CPM), one for the time course and one for the interaction (time course x pain). Nuisance regressors included in first-level GLMs were also included in the PPI models. The contrast for the psychological variable accounted for differences between CPM and control conditions, therefore testing for coupling changes between those two conditions. Contrast images were generated for the psycho-physiological interaction which tested for CPM-related changes in the correlation between regional time courses. A one-sample t test was calculated at the group level. Behavioural CPM effects (difference between CPM and control condition) were included as covariate in the second-level analyses. One participant had no significant voxel within the spinal seed region with our defined thresholds and thus could not be included into coupling analyses based on spinal time courses.

### Multiple comparisons correction

Correction for multiple comparisons of the functional imaging data in the brain was performed using different masks. For hypothesis testing that assumed changes in pain-related brain regions, a pain mask was used while for hypotheses that assumed changes in pain modulatory regions, a pain modulation mask was used. The pain mask was generated using different anatomical masks from the Neuromorphometrics atlas (Neuromorphometrics Inc., Somerville, MA, USA) such as the posterior and anterior insula, left postcentral gyrus, operculum, thalamus and mid-cingulate cortex. Moreover, a PAG mask (Faull et al., 2015) and a RVM mask (Brooks et al., 2017) were included to the pain mask (Fig. S10A). The RVM mask was smoothed by 1mm while the provided PAG mask is already smoothed by 2 mm. All masks were slightly warped as there were minor differences between different template brains and the final mask was cut to represent the group average field of view. The pain modulation mask included prefrontal regions such as the vmPFC and the perigenual ACC (pgACC), derived from Vega and colleagues (Vega et al., 2016) as well as the same PAG and RVM mask as previously mentioned (Fig. S10B). For spinal regions, we extracted the grey matter from the mean EPI image across all participants and manually cut the right dorsal horn. The resulting mask was adapted to spinal segment C6 based on the provided segments from the Spinal Cord Toolbox (Fig. S10C). Within these masks, we considered voxels significant at p < 0.05 using a family wise error (FWE) rate based on Gaussian random fields as implemented in SPM. Activation maps in main figures show only significant voxels within respective masks. Uncorrected whole-brain/spinal maps can be found in the data repository (see section ‘Data availability’). X, y and z coordinates are reported in MNI standard space.

### Preregistration and hypotheses

This study was preregistered at the German Clinical Trials Register (https://drks.de/search/en/trial/DRKS00027882). Our preregistered hypothesis for behavioural data states that pain ratings are decreased during the CPM condition compared to the control condition. Our main hypotheses for fMRI data says that BOLD responses are reduced in pain related brain regions in the CPM condition compared to control as well as in the spinal cord ipsilateral to stimulation site. Moreover, we hypothesized that CPM-related functional coupling between regions of the descending pain modulatory system is enhanced and that behavioural CPM magnitude correlates with BOLD reductions and coupling strength.

With regard to the preregistered ROIs and statistical correction for multiple comparisons correction, we deviated slightly from the original plan to be more consistent with our previous studies (Tinnermann et al., 2022, 2017). The preregistered ROIs do not differentiate between perception-related brain regions and modulation-related brain regions. As recent research has shown that regions such as the vmPFC/pgACC and PAG often show inverse activity patterns in relation to behavioural effects (Habermann and Büchel, 2025; Nold et al., 2025a; Tinnermann et al., 2022, 2017), we split the ROIs into two different masks for small volume correction. For the pain mask, we added some important pain related brain regions such as the opercular parts, the thalamus and the MCC and tested for CPM-related BOLD decreases within this mask. Within the pain modulation mask, we tested for CPM-related BOLD increases but did not consider the dorsolateral PFC as it is not formally part of the descending pain modulatory system and does not seem to modulate CPM (Ikarashi et al., 2025). Our spinal analyses were concentrated on spinal segment C6 as larger parts of C5 and C7 are missing in the final field of view due to motion correction and normalization procedures. Lastly, we did not use cluster-based correction methods as these have been heavily criticized in the past (Eklund et al., 2018, 2016; Noble et al., 2020; Woo et al., 2014) and hence we used hypothesis-driven peak voxel FWE corrections within our region of interest masks.

## Acknowledgements

The authors declare that they have no competing interests. All data needed to evaluate the conclusions in the paper are present in the paper and Supplementary Materials. Additional data such as second-level contrast images, behavioural data and masks for statistical comparison are provided on https://gin.g-node.org/atinnermann/cpm-corticospinal. Code for all analyses is openly available on GitHub: https://github.com/atinnermann/cpm-corticospinal.

AT, KEO, BH and CB were supported by the ERC AdG 883892 − PainPersist. AT was further supported by the German Research Foundation (TI 1110/1-1). CB and YC were supported by the SFB936 (936-178316478-A6).

We thank Kristian Hennings of Nocitech for his extensive help in setting up our experimental stimulation paradigm using the CPAR device. We also thank W. S. for his central role as the MR radiographer during our data acquisition.

## Notes

https://gin.g-node.org/atinnermann/cpm-corticospinal

### Competing Interest Statement

The authors have declared no competing interest.

