## Supplementary Figures for "Conditioned pain modulation recruits the descending pain modulatory system to inhibit spinal activity"

### Supplementary Material

#### Additional analyses

Behavioural control analysis

Since CPM effects can last beyond the duration of tonic stimulation, we assessed carry-over effects of CPM into control runs. Therefore, we compared participants that received CPM first and control second to participants that received control first and CPM second. In case that CPM effects would carry-over to the control condition, we should see a significant interaction between condition and condition order. However, this analysis was not significant ($\Delta VAS = 3.62$, $t_{27} = 0.79$, $p = 0.44$), indicating that CPM was not present during control conditions. Please note that because of our fully counterbalanced design, not all participants could be included in this analysis.

CPM effects assessed through HRF models

For reasons of completeness, we also analysed CPM effects with the traditional approach using HRF-convolved task regressors. This analysis yielded significant differences with lower activity in the CPM condition compared to the control condition in the right frontal operculum ($xyz_{MNI}$: 48/2/4, $t_{40} = 5.82$, $p_{SVC} = 0.007$) and left posterior insula and ($xyz_{MNI}$: -40/-15/0, $t_{40} = 5.12$, $p_{SVC} = 0.0499$; Fig. S8). The same analysis in the spinal cord did not yield any significant results (although a cluster in the right dorsal horn appeared at a very low statistical threshold; $xyz_{MNI}$ = 4/-48/-158, $t_{39} = 1.84$).

NPS and SIIPS responses in HRF models

In addition to time course analyses, we further tested the NPS and the Stimulus Intensity Independent Pain Signature (SIIPS) on traditional HRF models. The first analysis revealed a significantly lower NPS score for the CPM condition compared to the control condition ($\Delta NPS = 2.64$ au, $t_{40} = 3.47$, $p = 0.0013$; Fig. S9A). Similar to the time course analyses, we again compared the NPS scores separately for positive and negative weights, which confirm that most of the CPM-related differences in NPS scores stem from regions with a positive weight, i.e. pain-related brain regions (Fig. S9C). Similarly, SIIPS scores during CPM were significantly lower than during the control condition ($\Delta SIIPS = 503.75$ au, $t_{40} = 2.37$, $p = 0.023$; Fig. S9B).

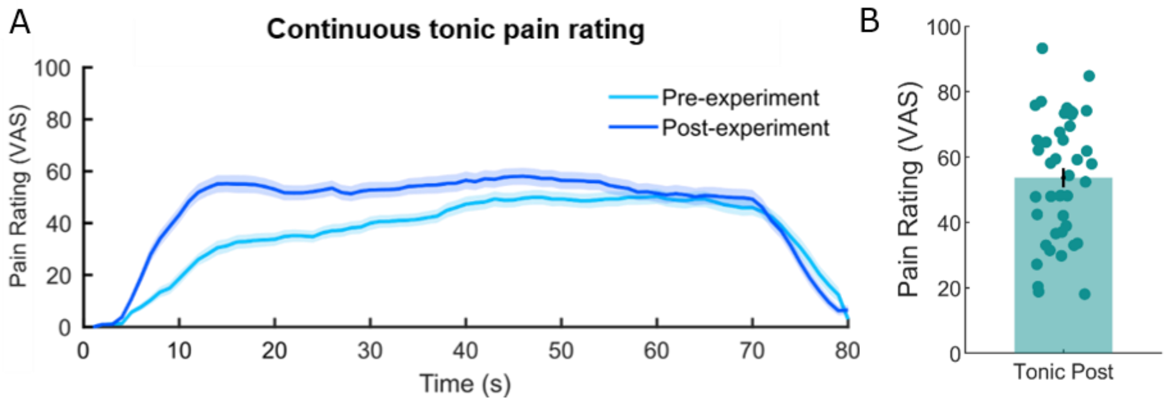


***Figure S1. Pre- and post-experiment continuous pain ratings for tonic pressure stimuli.***
*(**A**) Continuous pain rating for the tonic stimulus obtained right before and after the experiment. The stimulus was rated for 80 seconds. Sensitization to the conditioning stimulus over time was observed, with higher pain ratings in the first ~55 s of the stimulus in the post-experiment vs. pre-experiment rating session. (**B**) Distribution of averaged post ratings for the tonic stimulus excluding ramp times (first and last 10s).*

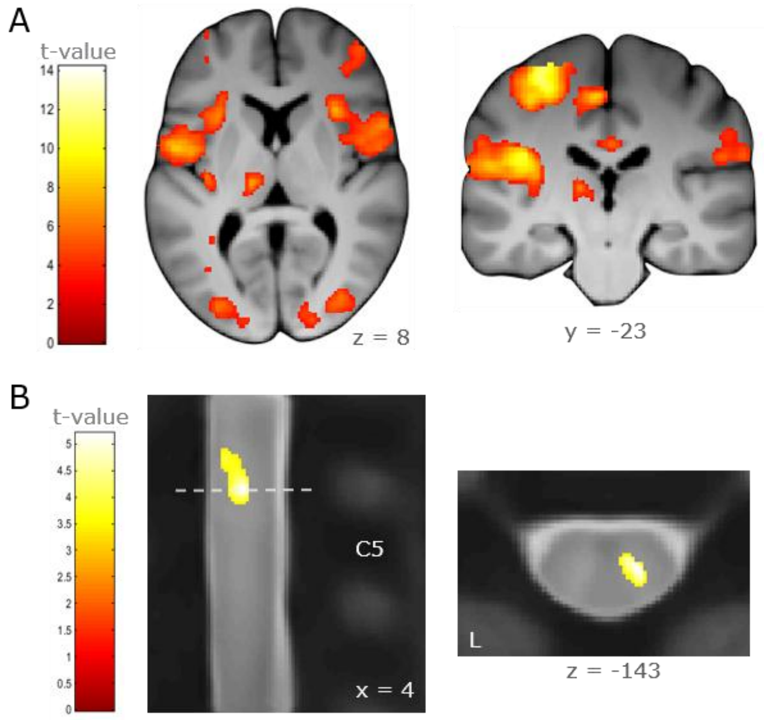


***Figure S2. Main effect of phasic pressure stimulation.*** *(**A**) Common pain-related brain regions show increased activation during phasic pressure stimulation (**B**) The right dorsal horn at the height of the cervical disk between C4 and C5 shows activity during phasic pressure stimulation. Uncorrected statistical t-maps are overlaid on an average structural T1 image in MNI template space for A) and dorsal horn region-of-interest masked t-maps are overlaid on a spinal mean EPI image in PAM50 template space for B). The visualization threshold is set to* $p_{uncorr} < 0.001$ *for the brain and spinal cord.*

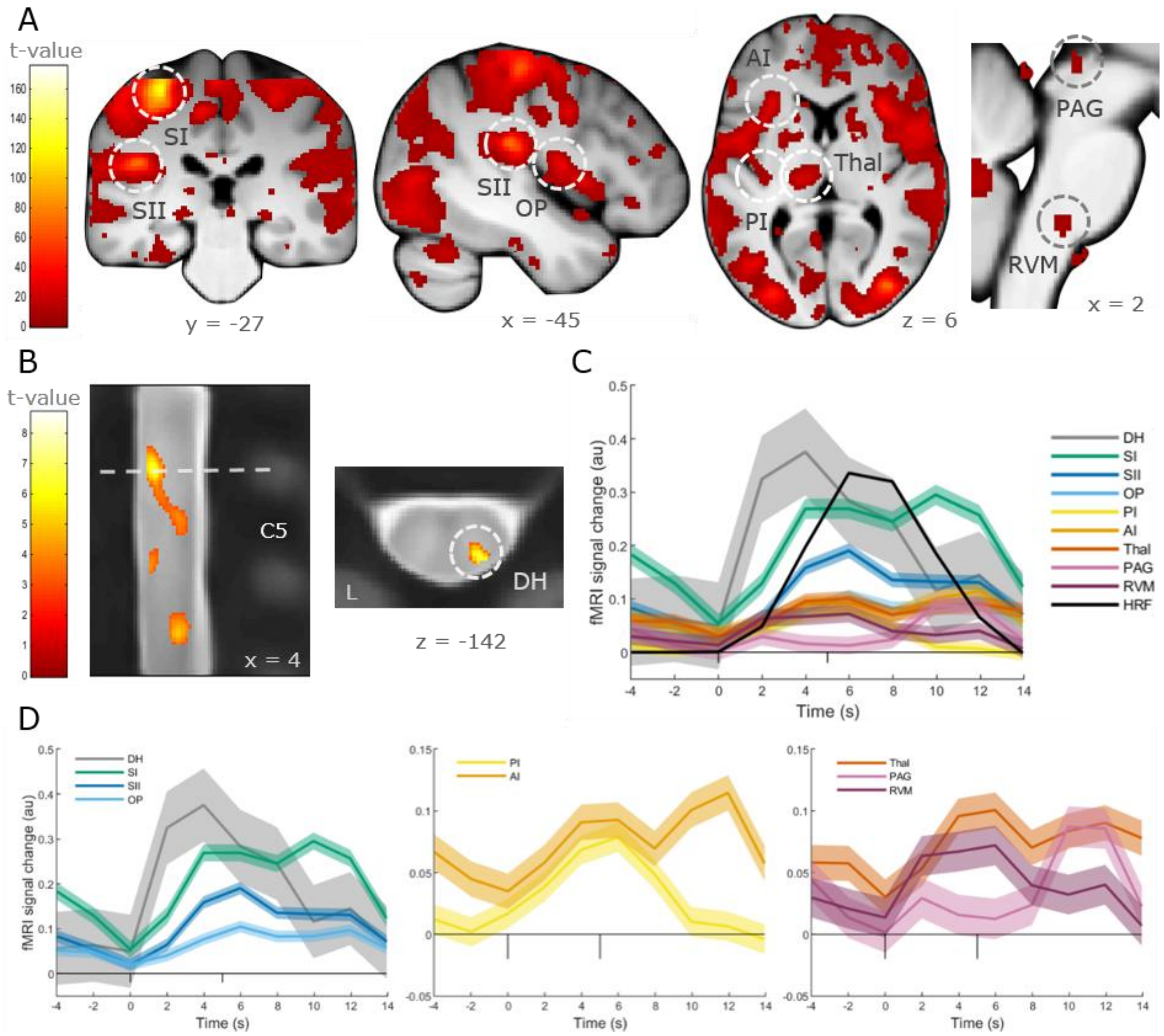


***Figure S3. Time courses of phasic pressure stimulation in the control condition.*** *(**A**) Effects-of-interest statistical F-maps are overlaid on an average structural T1 image in MNI template space and the visualization threshold is set to* $p_{uncorr} < 0.001$*. Dashed circles indicate regions from which time courses are plotted for the phasic pressure stimulation in the control condition. (**B**) Effects-of-interest statistical F-maps are overlaid on an average mean EPI image in PAM50 template space and the visualization threshold is set to* $p_{uncorr} < 0.001$*. Dashed circles indicate regions from which time courses are plotted for the phasic pressure stimulation in the control condition. (**C**) Time courses during phasic pressure stimulation for all pain-related regions. (**D**) The same time courses from C) are plotted in different graphs for better readability. Please note that the BOLD signal at a given time point reflects the averaged BOLD signal within one TR starting at that time point. DH: dorsal horn, SI: primary somatosensory cortex, SII: secondary somatosensory cortex, OP: operculum, PI: posterior insula, AI: interior insula, Thal: thalamus, PAG: periaqueductal gray, RVM: rostral ventromedial medulla, HRF: hemodynamic response function*

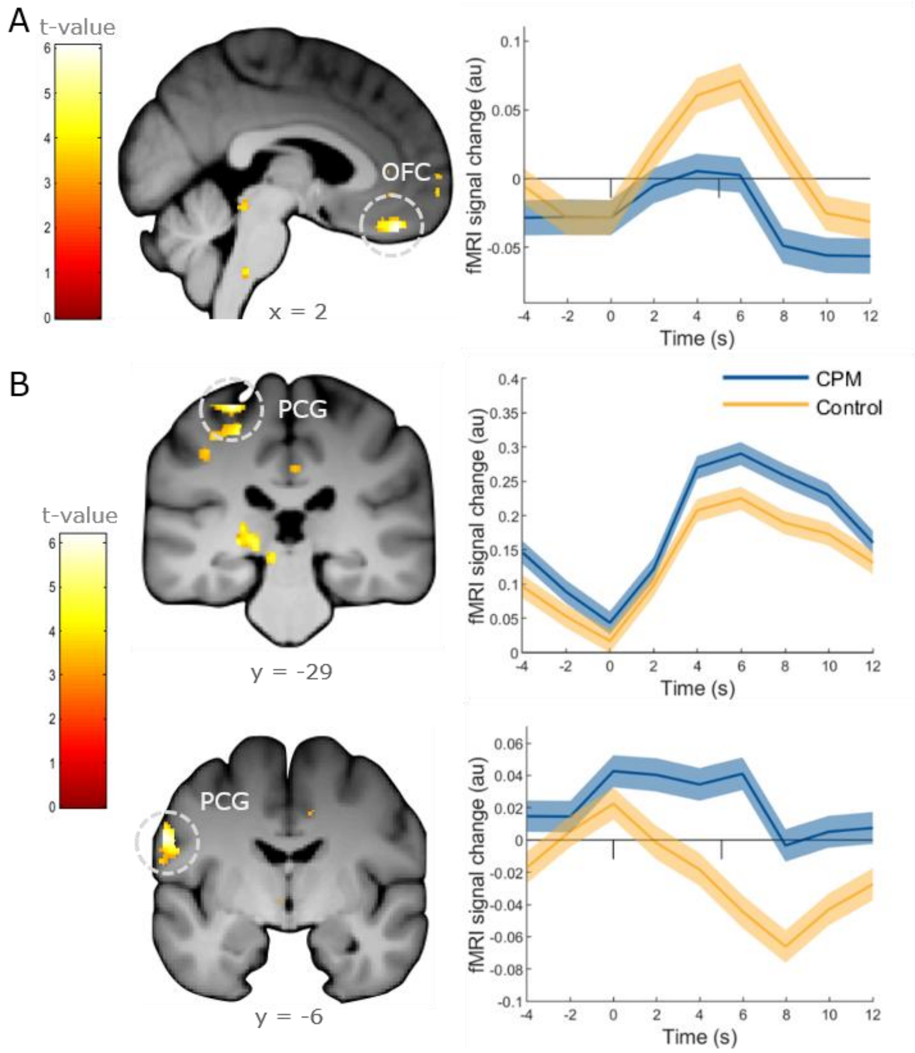


***Figure S4. Exploratory results.*** *(**A**) Decreased activation during the CPM condition within pain modulation regions was evident in the orbitofrontal cortex. (**B**) The postcentral gyrus showed increased activation during the CPM condition. A more superior region was activated in both, CPM and control condition, with a slightly higher activity during CPM while a more inferior region was activated during CPM and deactivated during the control condition. Masked uncorrected statistical t-maps are overlaid on an average structural T1 image in MNI template space and the visualization threshold is set to $p_{uncorr} < 0.001$. Please note that the BOLD signal at a given time point reflects the averaged BOLD signal within one TR starting at that time point.*

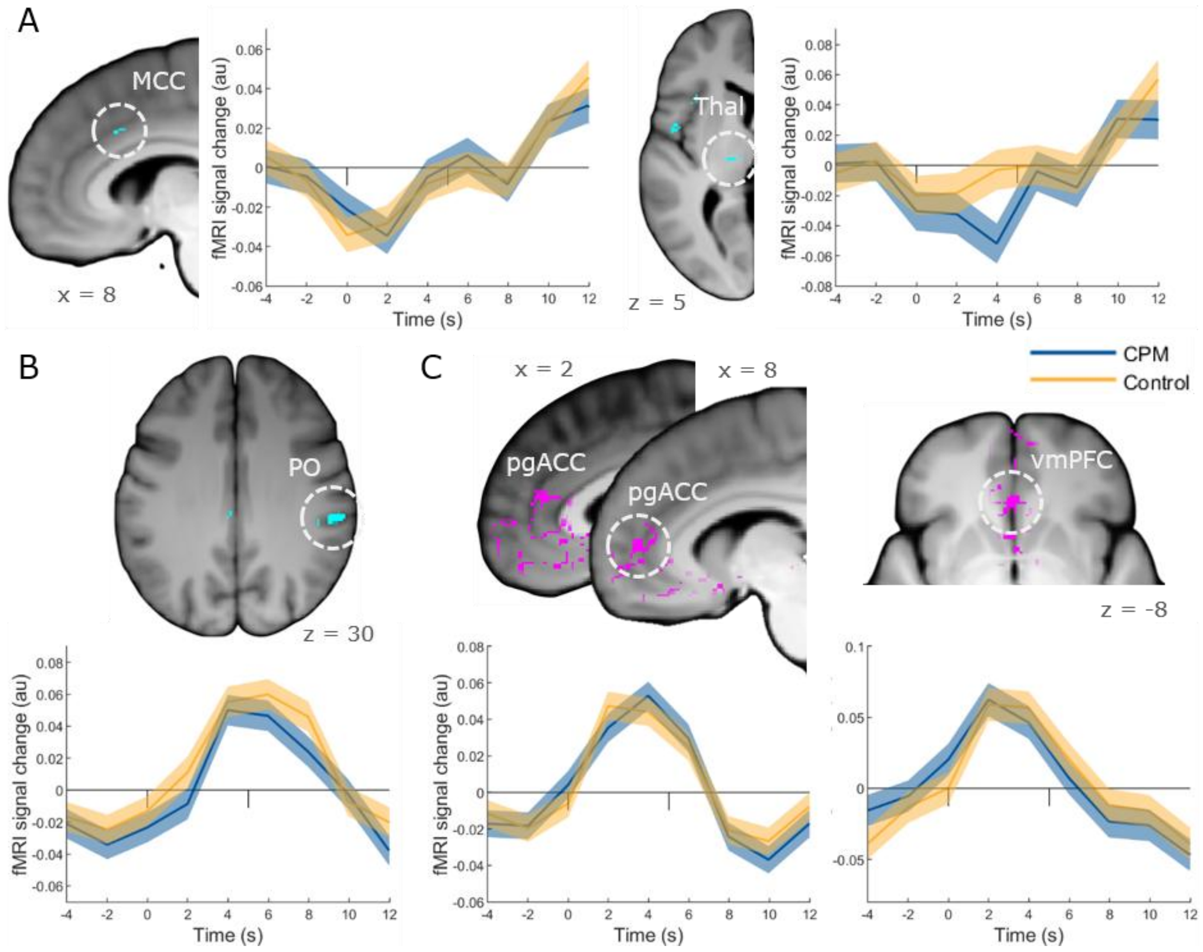


***Figure S5. Additional equivalence test results.*** *(**A**) Additional pain-related brain regions show equivalent responses for CPM and control such as the MCC* and thalamus*. (**B**) Responses ipsilateral to phasic stimulation are equivalent between CPM and control condition in the parietal operculum. (**C**) Time courses in pain-modulatory brain regions are equivalent between CPM and control conditions in the pgACC and vmPFC. Pain-related (A and B) and pain-modulatory (C) region-of-interest masked binary equivalence maps representing voxels that meet a defined equivalence criterion are overlaid on an average structural T1 image in MNI template space. Please note that the BOLD signal at a given time point reflects the averaged BOLD signal within one TR starting at that time point. MCC: mid cingulate cortex, Thal: thalamus, PO: parietal operculum, pgACC: perigenual anterior cingulate cortex, vmPFC: ventromedial prefrontal cortex*

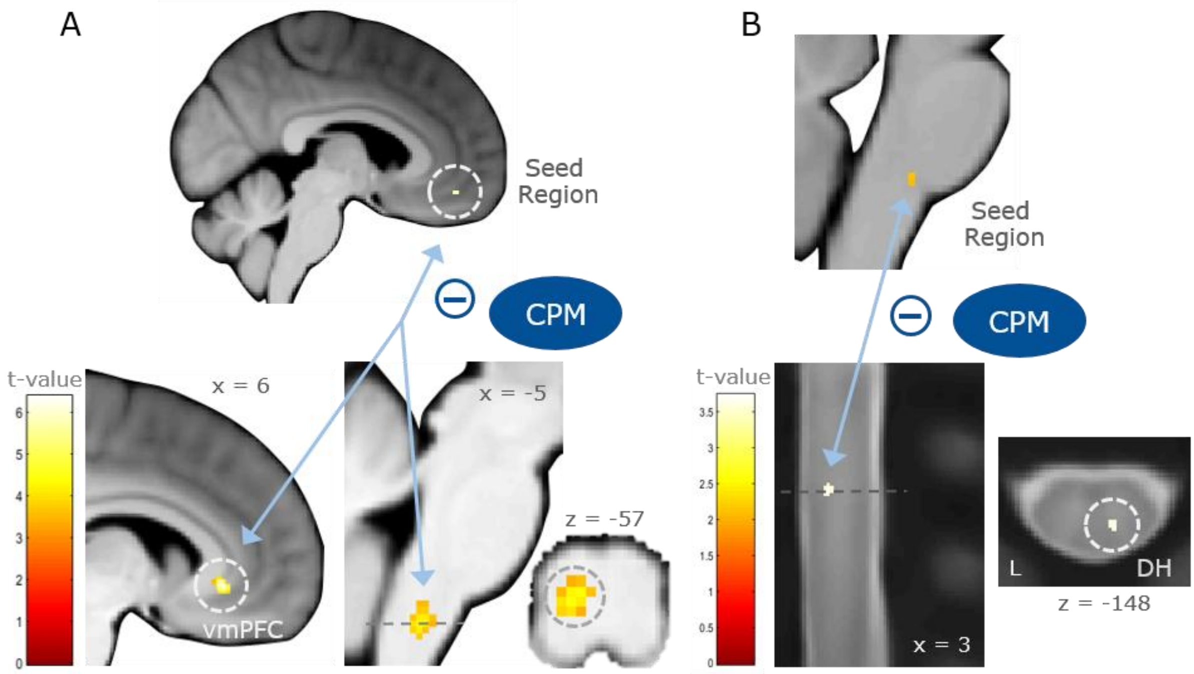


***Figure S6. Additional CPM-related coupling results along regions of the descending pain modulatory pathway.** (**A**) Coupling strength between the vmPFC and another vmPFC subregion was decreased during CPM. An additional cluster appeared in the brainstem, supposedly the subnucleus dorsalis reticularis, which was not part of our hypothesis-driven analysis. (**B**) Coupling strength between the RVM and right dorsal horn of spinal segment C6 was decreased under CPM. The visualization threshold is set to $p < 0.001$ for A) and $p_{SVC} < 0.05$ for B). Please note that the BOLD signal at a given time point reflects the averaged BOLD signal within one TR starting at that time point. vmPFC: ventromedial prefrontal cortex, DH: dorsal horn*

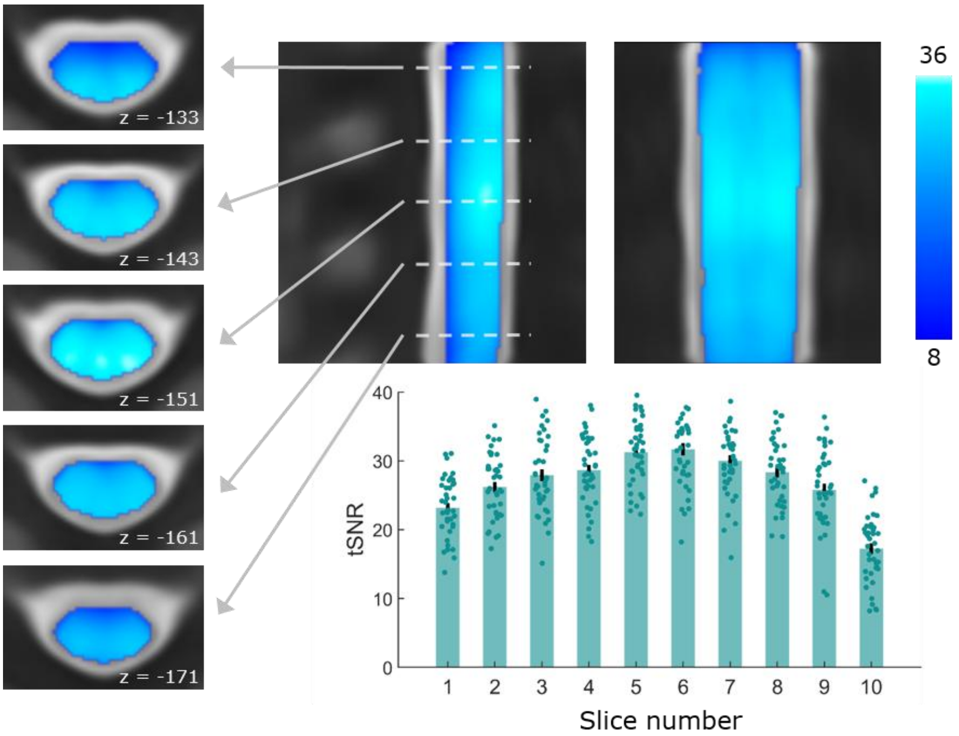


***Figure S7. Temporal signal-to-noise ratios in the spinal cord.*** *tSNR maps were calculated after motion correction, subsequently warped to PAM50 template space and averaged across participants. The bar graphs depict tSNR per slice in native space and averaged across participants. Dots depict tSNR values from individual participants.*

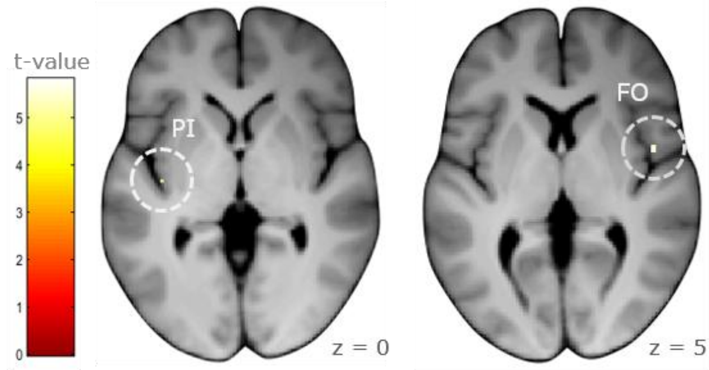


***Figure S8. CPM effects in a standard GLM analysis.*** *Significant activation differences between control and CPM conditions in a standard GLM analysis were localized in the right operculum and left posterior insula. Pain-related region-of-interest masked statistical t-maps are overlaid on an average structural T1 image in MNI template space and the visualization threshold is set to* $p_{SVC} < 0.05$.

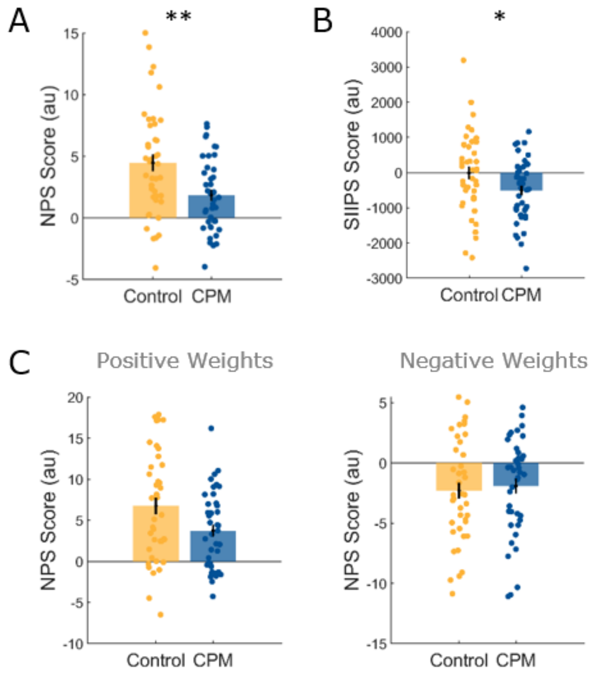


***Figure S9. Neurological Pain Signature and Stimulus Intensity Independent Pain Signature responses in HRF models.*** *(**A**) NPS scores for CPM and control condition show a significant lower score during CPM. (**B**) SIIPS scores for CPM and control condition show a significant lower score during CPM. (**C**) NPS scores analysed separately for positive and negative weights show that most of the difference of NPS response is related to regions with a positive weight.*

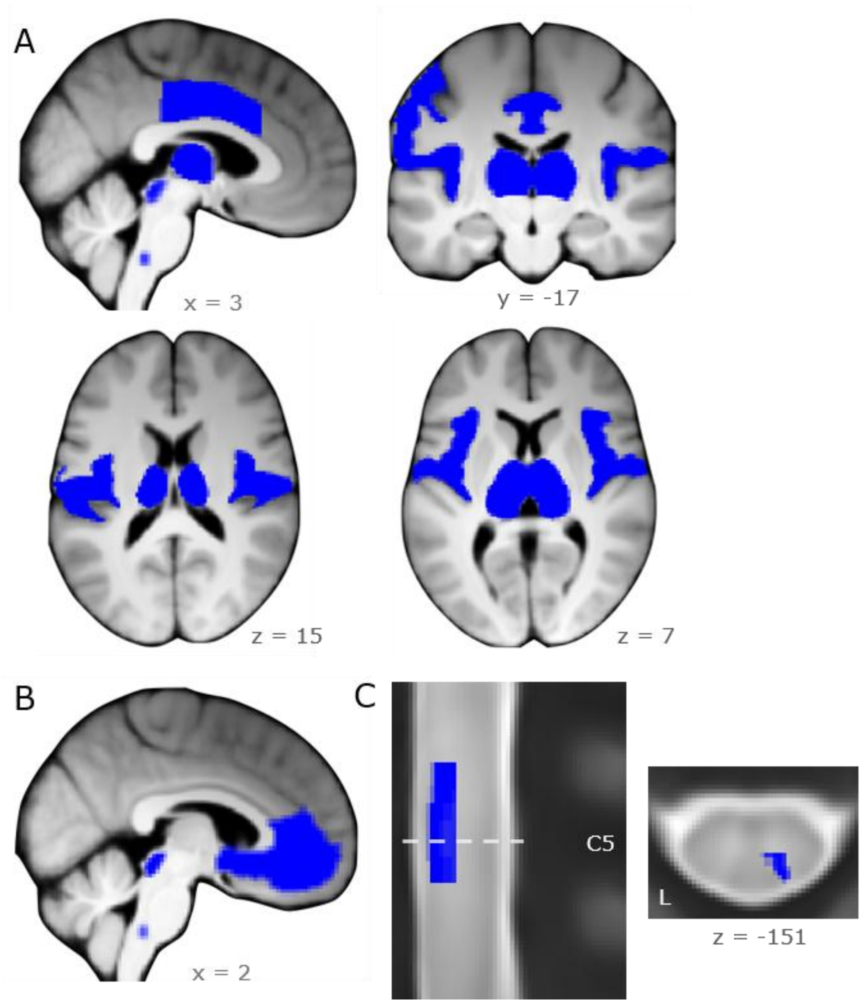


***Figure S10. Masks for small volume correction.*** *(**A**) Combined anatomical masks from the Neuromorphometrics atlas that cover pain-related brain regions such as opercular and insular regions, mid cingulate cortex, thalamus, left postcentral gyrus, PAG and RVM. (**B**) Pain modulation mask covering the vmPFC, PAG and RVM. (**C**) Right dorsal horn mask for spinal segment C6.*
